# Flexible Experimental Designs for Valid Single-cell RNA-sequencing Experiments Allowing Batch Effects Correction

**DOI:** 10.1101/533372

**Authors:** Fangda Song, Ga Ming Angus Chan, Yingying Wei

## Abstract

Despite their widespread applications, single-cell RNA-sequencing (scRNA-seq) experiments are still plagued by batch effects and dropout events. Although the completely randomized experimental design has frequently been advocated to control for batch effects, it is rarely implemented in real applications due to time and budget constraints. Here, we mathematically prove that under two more flexible and realistic experimental designs—the “reference panel” and the “chain-type” designs—true biological variability can also be separated from batch effects. We develop **B**atch effects correction with **U**nknown **S**ubtypes for scRNA-**seq** data (BUSseq), which is an interpretable Bayesian hierarchical model that closely follows the data-generating mechanism of scRNA-seq experiments. BUSseq can simultaneously correct batch effects, cluster cell types, impute missing data caused by dropout events, and detect differentially expressed genes without requiring a preliminary normalization step. We demonstrate that BUSseq outperforms existing methods with simulated and real data.

## Introduction

Single-cell RNA-sequencing (scRNA-seq) technologies enable the measurement of the transcriptome of individual cells, which provides unprecedented opportunities to discover cell types and understand cellular heterogeneity [1]. However, like the other high-throughput technologies [2–4], scRNA-seq experiments can suffer from severe batch effects [5]. Moreover, compared to bulk RNA-seq data, scRNA-seq data can have an excessive number of zeros that result from dropout events—that is, the expressions of some genes are not detected even though they are actually expressed in the cell due to amplification failure prior to sequencing [6]. Consequently, despite the widespread adoption of scRNA-seq experiments, the design of a valid scRNA-seq experiment that allows the batch effects to be removed, the biological cell types to be discovered, and the missing data to be imputed remains an open problem.

One of the major tasks of scRNA-seq experiments is to identify cell types for a population of cells [1]. The cell type of each individual cell is unknown and is often the target of inference. Classic batch effects correction methods, such as Combat [7] and SVA [8, 9], are designed for bulk experiments and require knowledge of the subtype information of each sample a prior. For scRNA-seq data, this subtype information corresponds to the cell type of each individual cell. Clearly, these methods are thus infeasible for scRNA-seq data. Alternatively, if one has knowledge of a set of control genes whose expression levels are constant across cell types, then it is possible to apply RUV [10, 11]. However, selecting control genes is often difficult for scRNA-seq experiments.

To identify unknown subtypes, MetaSparseKmeans [12] jointly clusters samples across batches. Unfortunately, MetaSparseKmeans requires all subtypes to be present in each batch. Suppose that we conduct scRNA-seq experiments for blood samples from a healthy individual and a leukemia patient, one person per batch. Although we can anticipate that the two batches will share T cells and B cells, we do not expect that the healthy individual will have cancer cells as the leukemia patient. Therefore, MetaSparseKmeans is too restrictive for many scRNA-seq experiments.

The mutual-nearest-neighbor (MNN) based approaches, including MNN [13] and Scanorama [14], allow each batch to contain some but not all cell types. However, these methods require batch effects to be almost orthogonal to the biological subspaces and much smaller than the biological variations between different cell types [13]. These are strong assumptions and cannot be validated at the design stage of the experiments. Seurat [15, 16], LIGER [17] and scMerge [18] attempt to identify shared variations across batches by low-dimensional embeddings and treat them as shared cell types. However, they may mistake the technical artifacts as the biological variability of interest if some batches share certain technical noises, for example when each patient is measured by several batches. To handle severe batch effects for microarray data, Luo and Wei [19] developed BUS to simultaneously cluster samples across multiple batches and correct batch effects. However, none of the above methods considers features unique to scRNA-seq data, such as the count nature of the data, over-dispersion [20], dropout events [6], or cell-specific size factors [21]. ZIFA [22] and ZINB-WaVE [23] incorporate dropout events into the factor model, whereas scVI [24] and SAVER-X [25] couple the modeling of dropout events with neural networks. However, as is the case with the other state-of-the-art methods, these papers do not discuss the designs of scRNA-seq experiments under which their methods are applicable.

Nevertheless, it is crucial to understand the conditions under which biological variability can be separated from technical artifacts. Obviously, for completely confounded designs—for example one in which batch 1 measures cell type 1 and 2, whereas batch 2 measures cell type 3 and 4—no method is applicable.

Here, we propose Batch effects correction with Unknown Subtypes for scRNA-seq data (BUSseq), an interpretable hierarchical model that simultaneously corrects batch effects, clusters cell types, and takes care of the count data nature, the overdispersion, the dropout events, and the cell-specific size factors of scRNA-seq data. We mathematically prove that it is legitimate to conduct scRNA-seq experiments under not only the commonly advocated completely randomized design [1, 5, 26, 27], in which each batch measures all cell types, but also the “reference panel” design and the “chain-type” design, which allow some cell types to be missing from some batches. Furthermore, we demonstrate that BUSseq outperforms the existing approaches in both simulation data and real applications. The theoretical results answer the question about when we can integrate multiple scRNA-seq datasets and analyze them jointly. We envision that the proposed experimental designs will be able to guide biomedical researchers and help them to design better scRNA-seq experiments.

## Results

### BUSseq is an interpretable hierarchical model for scRNA-seq

We develop a hierarchical model BUSseq that closely mimics the data generating procedure of scRNA-seq experiments (**Fig. 1a** and **Supplementary Fig. 1**). Given that we have measured *B* batches of cells each with a sample size of *n*_*b*_, let us denote the underlying gene expression level of gene *g* in cell *i* of batch *b* as *X*_*big*_. *X*_*big*_ follows a negative binomial distribution with mean expression level *µ*_*big*_ and a gene-specific and batch-specific overdispersion parameter *ϕ*_*bg*_. The mean expression level is determined by the cell type *W*_*bi*_ with the cell type effect *β*_*gk*_, the log-scale baseline expression level *α*_*g*_, the location batch effect *ν*_*bg*_, and the cell-specific size factor *δ*_*bi*_. The cell-specific size factor *δ*_*bi*_ characterizes the impact of cell size, library size and sequencing depth. It is of note that the cell type *W*_*bi*_ of each individual cell is unknown and is our target of inference. Therefore, we assume that a cell on batch *b* comes from cell type *k* with probability *P* (*W*_*bi*_ = *k*) = *π*_*bk*_ and the proportions of cell types (*π*_*b*1_, …, *π*_*bK*_) vary among batches.

**Figure 1:**
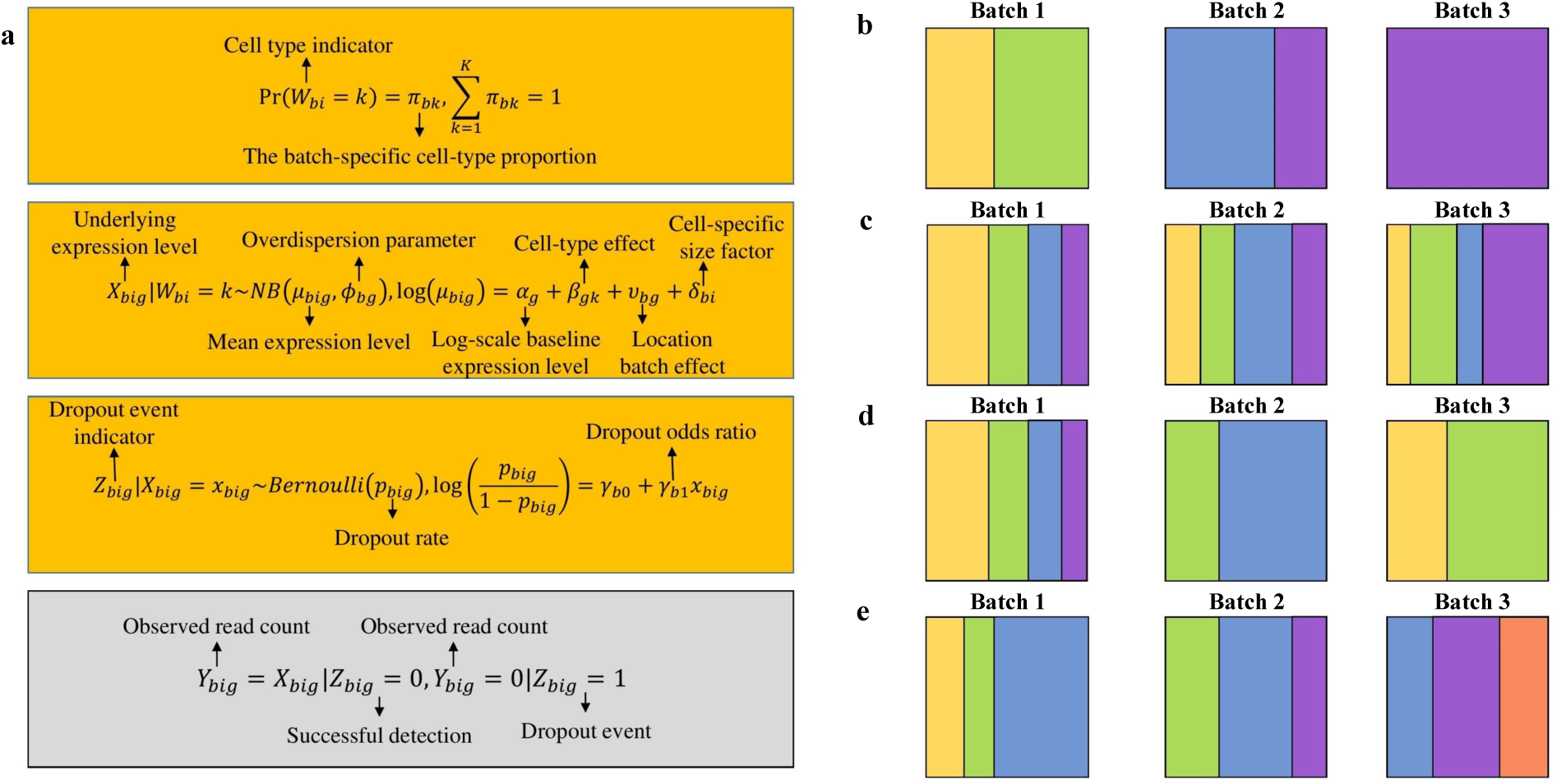
Illustration of the BUSseq model and various types of experimental designs. **(a)** The hierarchical structure of the BUSseq model. Only *Y*_*big*_ in the grey rectangle is observed. **(b)** A confounded design that contains three batches. Each polychrome rectangle represents one batch of scRNA-seq data with genes in rows and cells in columns; and each color indicates a cell type. Batch 1 assays cells from cell types 1 and 2; batch 2 profiles cells from cell types 3 and 4; and batch 3 only contains cells from cell type 4. **(c)** The complete setting design. Each batch assays cells from all of the four cell types, although the cellular compositions vary across batches. **(d)** The reference panel design. Batch 1 contains cells from all of the cell types, and all the other batches have at least two cell types. **(e)** The chain-type design. Every two consecutive batches share two cell types. Batch 1 and Batch 2 share cell types 2 and 3; Batch 2 and Batch 3 share cell types 3 and 4 (see also **Supplementary Figs. 1 and 2**).

Unfortunately, it is not always possible to observe the expression level *X*_*big*_. Without dropout (*Z*_*big*_ = 0), we can directly observe *Y*_*big*_ = *X*_*big*_. However, if a dropout event occurs (*Z*_*big*_ = 1), then we observe *Y*_*big*_ = 0 instead of *X*_*big*_. It has been noted that highly expressed genes are less-likely to suffer from dropout events [6]. We thus model the dependence of the dropout rate *P* (*Z*_*big*_ = 1|*X*_*big*_) on the expression level using a logistic regression with batch-specific intercept *γ*_*b*0_ and odds ratio *γ*_*b*1_.

Noteworthy, BUSseq includes the negative binomial distribution without zero inflation as a special case. When all cells are from a single cell type and the cell-specific size factor *δ*_*bi*_ is estimated a priori according to spike-in genes, BUSseq can reduce to a form similar to BASiCS [20].

We only observe *Y*_*big*_ for all cells in the *B* batches and the total *G* genes. We conduct statistical inference under the Bayesian framework and develop a Markov chain Monte Carlo (MCMC) algorithm [28]. Based on the parameter estimates, we can learn the cell type for each individual cell, impute the missing underlying expression levels *X*_*big*_ for dropout events, and identify genes that are differentially expressed among cell types. Moreover, our algorithm can automatically detect the total number of cell types *K* that exists in the dataset according to the Bayesian information criterion (BIC) [29]. BUSseq also provides a batch-effect corrected version of count data, which can be used for downstream analysis as if all of the data were measured in a single batch. Details are in Methods and Supplementary Notes.

### Valid experimental designs for scRNA-seq experiments

If a study design is completely confounded, as shown in **Fig. 1b**, then no method can separate biological variability from technical artifacts, because different combinations of batch-effect and cell-type-effect values can lead to the same probabilistic distribution for the observed data, which in statistics is termed a *non-identifiable* model. Formally, a model is said to be *identifiable* if each probability distribution can arise from only one set of parameter values [30]. Statistical inference is impossible for non-identifiable models because two sets of distinct parameter values can give rise to the same probabilistic function. We prove that the BUSseq model is identifiable under conditions that are very easily met in reality. It is thus applicable to a wide range of experimental designs.

For the “complete setting,” in which each batch measures all of the cell types (**Fig. 1c** and Theorem 1 in Methods), BUSseq is identifiable as long as: (I) the odds ratio *γ*_*b*1_s in the logistic regressions for the dropout rates are negative for all of the batches, (II) every two cell types have more than one differentially expressed gene, and (III) the ratios of mean expression levels between two cell types 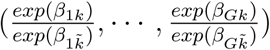 are different for each cell-type pair 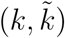 (see Theorem 1 in Methods). Condition (I) requires that the highly expressed genes are less likely to have dropout events, which is routinely observed for scRNA-seq data [6]. Condition (II) always holds in reality. Because scRNA-seq experiments measure the whole transcriptome of a cell, condition (III) is also always met in real data. For example, if there exists one gene *g* such that for any two distinct cell-type pairs (*k*_1_, *k*_2_) and (*k*_3_, *k*_4_) their mean expression levels ratios 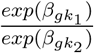 and 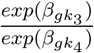 are not the same, then condition (III) is already satisfied.

The commonly advocated completely randomized experimental design falls into the “complete setting” design, whereas the latter further relaxes the assumption implied by the former that the cell-type proportions are almost the same for all batches. The identical composition of the cell population within each batch is a crucial requirement for traditional batch effects correction methods developed for bulk experiments such as Combat [13]. In contrast, BUSseq is not limited to this balanced design constraint and is applicable to not only the completely randomized design but also the general complete setting design.

Ideally, we would wish to adopt completely randomized experimental designs. However, in reality, it is always very challenging to implement complete randomization due to time and budget constraints. For example, when we recruit patients sequentially, we often have to conduct scRNA-seq experiments patient-by-patient rather than randomize the cells from all of the patients to each batch, and the patients may not have the same set of cell types. Fortunately, we can prove that BUSseq also applies to two sets of flexible experimental designs, which allow cell types to be measured in only some but not all of the batches.

Assuming that conditions (I)-(III) are satisfied, if there exists one batch that contains cells from all cell types and the other batches have at least two cell types (**Fig. 1d**), then BUSseq can tease out the batch effects and identify the true biological variability (see Theorem 2 in Methods). We call this setting the “reference panel design.”

Sometimes, it can still be difficult to obtain a reference batch that collects all cell types. In this case, we can turn to the chain-type design, which requires every two consecutive batches to share two cell types (**Fig. 1e**). Under the chain-type design, given that conditions (I)-(III) hold, BUSseq is also identifiable and can estimate the parameters well (see Theorem 3 in Methods).

A special case of the chain-type design is when two common cell types are shared by all of the batches, which is frequently encountered in real applications. For instance, when blood samples are assayed, even if we perform scRNA-seq experiment patient-by-patient with one patient per batch, we know a priori that each batch will contain at least both T cells and B cells, thus satisfying the requirement of the chain-type design.

The key insight is that despite batch effects, differences between cell types remain constant across batches. The differences between a pair of cell types allow us to distinguish batch effects from biological variability for those batches that measure both cell types. Once batch effects have been identified, we can conduct joint clustering across batches with batch effects adjusted. In fact, BUSseq can separate batch effects from cell type effects under more general designs beyond the easily understood and commonly encountered reference panel design and chain-type design. If we regard each batch as a node in a graph and connect two nodes with an edge if the two batches share at least two cell types, then BUSseq is identifiable as long as the resulting graph is connected (see **Supplementary Fig. 2** and Theorem 4 in Methods).

For scRNA-seq data, dropout rate depends on the underlying expression levels. Such missing data mechanism is called missing not at random (MNAR) in statistics. It is very challenging to establish identifiability for MNAR. Miao et al. [31] showed that for many cases even when both the outcome distribution and the missing data mechanism have parametric forms, the model can be nonidentifiable. However, fortunately, despite the dropout events and the cell-specific size factors, by creating a set of functions similar to the probability generating function, we proved Theorems 1–4 (see their proofs in Supplementary Notes). The reference panel design, the chain-type design and the connected design liberalize researchers from the ideal but often unrealistic requirement of the completely randomized design.

### BUSseq accurately estimates the parameters and imputes the missing data

We first evaluate the performance of BUSseq via a simulation study. We simulate a dataset with four batches and a total of five cell types under the chain-type design (**Fig. 2a-d** and Theorem 3). Every two consecutive batches share at least two cell types, but none of the batches contains all of the cell types. The sample sizes for each batch are (*n*_1_, *n*_2_, *n*_3_, *n*_4_) = (300, 300, 200, 200), and there are a total of 3,000 genes. In real datasets, batch effects are often much larger than the cell type effects (**Fig. 3a**) and not orthogonal to the cell type effects (**Supplementary Fig. 3**). In the simulation study, we choose the magnitude of the batch effects, cell type effects, the dropout rates, and the cell-specific size factors to mimic real data scenarios (**Fig. 3a**). The simulated observed data suffer from severe batch effects and dropout events (**Fig. 2d** and **Fig. 3c**). The dropout rates for the four batches are 26.79%, 24.53%, 28.36% and 31.29%, with the corresponding total zero proportions given by 44.13%, 48.85%, 53.07% and 61.38%.

**Figure 2:**
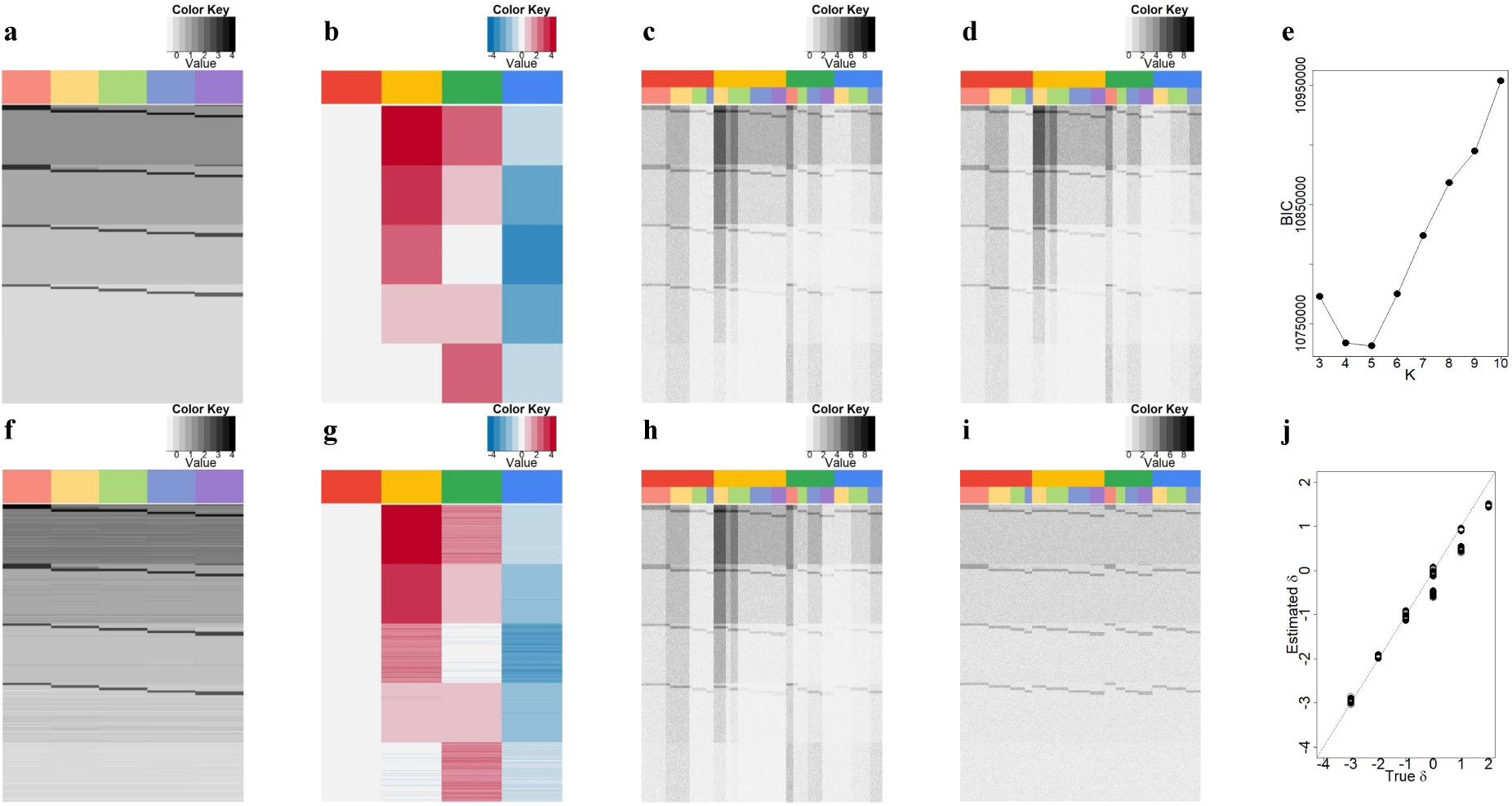
Patterns of the simulation study. **(a)** True log-scale mean expression levels for each cell type ***α*** + ***β***. Each row represents a gene, and each column corresponds to a cell type. The intrinsic genes that are differentially expressed between cell types can have high, medium high, median low or low expression levels. **(b)** True batch effects. Each row represents a gene, and each column corresponds to a batch. **(c)** True underlying expression levels ***X***. Each row represents a gene, and each column corresponds to a cell. The upper color bar indicates the batches, and the lower color bar represents the cell types. There are a total of 3,000 genes. The sample sizes for each batch are 300, 300, 200 and 200, respectively. **(d)** The simulated observed data ***Y***. The overall dropout rate is 27.3%, whereas the overall zero rate is 50.8%. **(e)** The BIC plot. The BIC attains the minimum at *K* = 5, identifying the true cell type number. **(f)** The estimated log-scale mean expression levels for each cell type 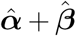.**(g)** Estimated batch effects. **(h)** Imputed expression levels 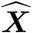. **(i)** Corrected count data 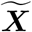 grouped by batches. **(j)** Scatter plot of the estimated versus the true cell-specific size factors.

**Figure 3:**
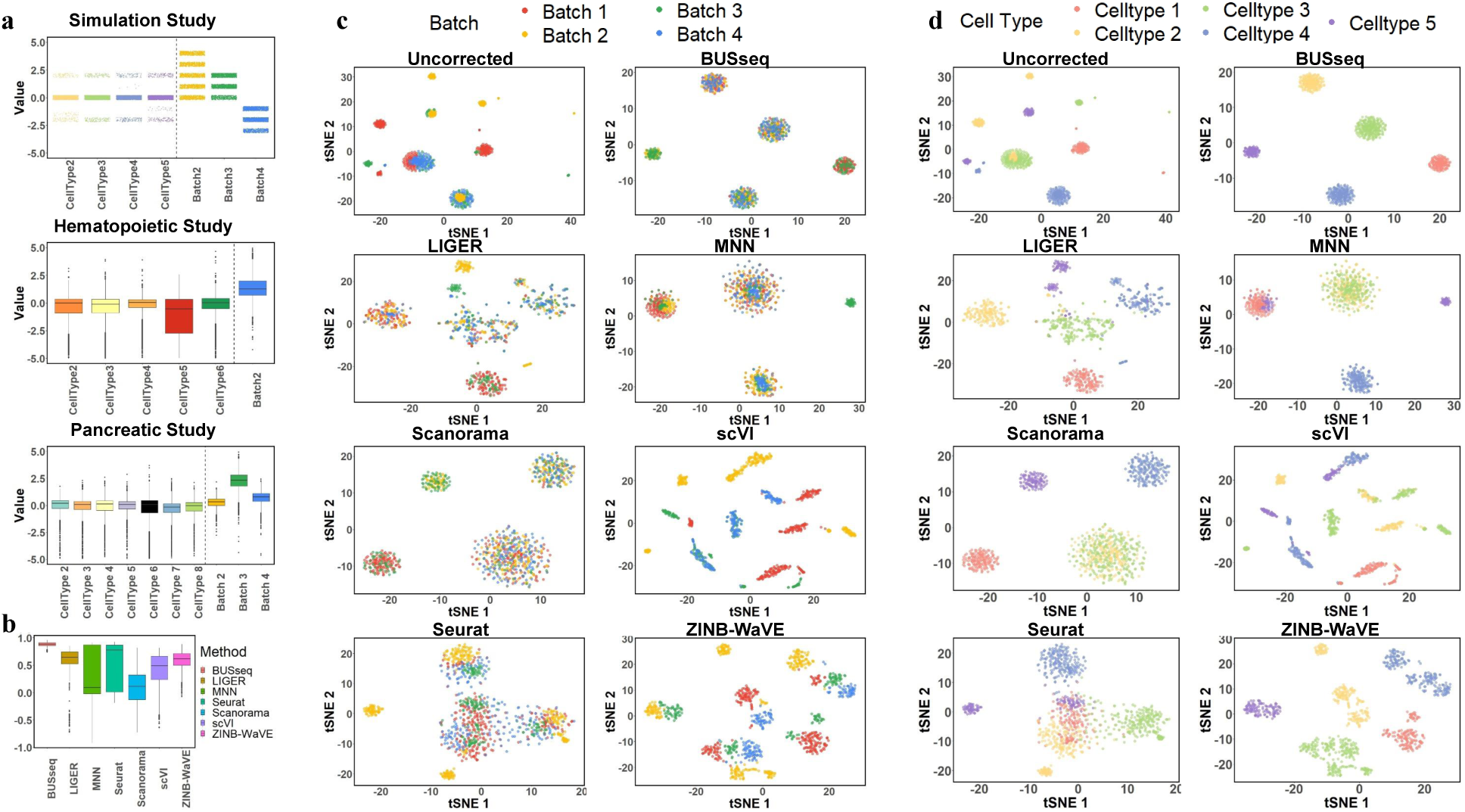
Comparison of batch effects correction methods in the simulation study. **(a)** Comparison of the magnitude of cell type effects and batch effects in the simulation study and the two real applications. The subpanel for the simulation study jitters around the assumed values for ***β*** and ***ν***. The boxplots show the distributions of the estimated cell type effects 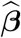 and batch effects 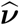 by BUSseq in the two real studies. The magnitude of the batch effects and cell type effects in the simulation study were chosen to mimic the real data scenarios. **(b)** The boxplots of silhouette coefficients for all compared methods. **(c)** T-distributed Stochastic Neighbor Embedding (t-SNE) plots colored by batch for each compared method. **(d)** t-SNE plots colored by true cell type labels for each compared method.

BUSseq correctly identifies the presence of five cell types among the cells (**Fig. 2e**). Moreover, despite the dropout events, BUSseq accurately estimates the cell type effects *β*_*gk*_s (**Fig. 2a and f**), the batch effects *ν*_*bg*_s (**Fig. 2b and g**), and the cell-specific size factors *δ*_*bi*_s (**Fig. 2j**). When controlling the Bayesian False Discovery Rate (FDR) at 0.05 [32, 33], we identify all intrinsic genes that differentiate cell types with the true FDR being 0.02 (Methods).

In the simulation study, we know the underlying expression levels *X*_*big*_s. Therefore, we can compare them with our inferred expression levels 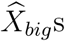 based the observed data *Y*_*big*_s which are subject to dropout events. **Fig. 2h** demonstrates that BUSseq can learn the underlying expression levels well. This success arises because BUSseq uses an integrative model to borrow strengths both across genes and across cells from all batches. As a result, BUSseq can achieve accurate estimation and imputation despite the dropout events.

Combat offers a version of data that have been adjusted for batch effects [7]. Here, we also provide batch-effects-corrected count data based on quantile matching (Methods). The adjusted count data no longer suffer from batch effects and dropout events, and they even do not need further cell-specific normalization (Fig. **2i**). Therefore, they can be treated as if measured in a single batch for downstream analysis.

### BUSseq outperforms existing methods in simulation study

We benchmarked BUSseq with the state-of-the-art methods for batch effects correction for scRNA-seq data—LIGER [17], MNN [13], Scanorama [14], scVI [24], Seurat [16] and ZINB-WaVE [23]. The adjusted Rand index (ARI) measures the consistency between two clustering results and is between zero and one, a higher value indicating better consistency (Supplementary Notes). The ARI between the inferred cell types 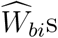 by BUSseq and the true underlying cell types *W*_*bi*_s is one. Thus, BUSseq can perfectly recover the true cell type of each cell. In comparison, we apply each of the compared methods to the dataset and then perform their own clustering approaches (Supplementary Notes). The ARI is able to compare the consistency of two clustering results even if the numbers of clusters differ, therefore, we choose the number of cell types by the default approach of each method rather than set it to a common number. The resulting ARIs are 0.837 for LIGER, 0.654 for MNN, 0.521 for Scanorama, 0.480 for scVI, 0.632 for Seurat and 0.571 for ZINB-WaVE. Moreover, the t-SNE plots (**Fig. 3c and d**) show that only BUSseq can perfectly cluster the cells by cell types rather than batches. We also calculated the silhouette score for each cell for each compared method (Supplementary Notes). A high silhouette score indicates that the cell is well matched to its own cluster and separated from neighboring clusters. **Fig. 3b** shows that BUSseq gives the best segregated clusters.

### BUSseq outperforms existing methods on hematopoietic data

We re-analyzed the two hematopoietic datasets previously studied by Haghverdi et al.[13], one profiled by the SMART-seq2 protocol for a population of hematopoietic stem and progenitor cells (HSPC) from 12-week-old female mice [34] and another assayed by the massively parallel single-cell RNA-sequencing (MARS-seq) protocol for myeloid progenitors from 6-to 8-week-old female mice [35]. Although the two datasets were generated in two different laboratories (**Fig. 4a**), both datasets have cell-type label for each cell that is annotated according to the expression levels of marker genes [13, 35] from fluorescence-activated cell sorting (FACS) (Methods).

**Figure 4:**
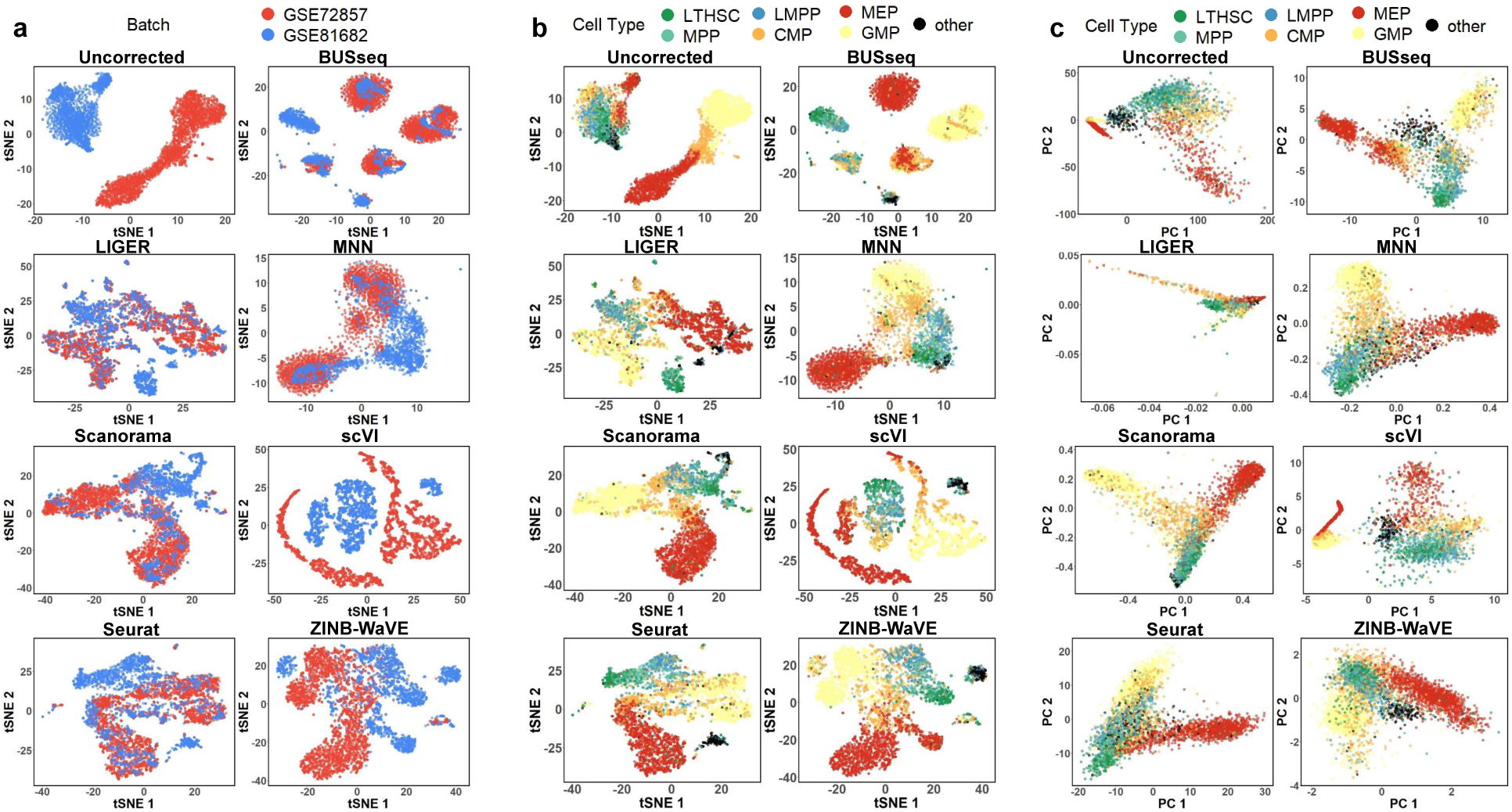
t-SNE and Principal Component Analysis (PCA) plots for the hematopoietic data. **(a)** t-SNE plots colored by batch. **(b)** t-SNE plots colored by FACS cell type labels. (c) PCA plots colored by FACS cell type labels.

In order to compare BUSseq with existing methods, we compute the ARI between the clustering of each method and the FACS labels. The resulting ARIs are 0.582 for BUSseq, 0.307 for LIGER, 0.575 for MNN, 0.518 for Scanorama, 0.197 for scVI, 0.266 for Seurat and 0.348 for ZINB-WaVE. BUSseq thus outperforms all of the other methods in being consistent with FACS labeling. BUSseq also has silhouette coefficients that are comparable to those of MNN, which are better than those of all the other methods (**Supplementary Fig. 4**). Furthermore, t-SNE plots confirm that BUSseq performs the best in segregating cells into different cell types (**Fig. 4b**).

Specifically, BUSseq learns 6 cell types from the dataset. According to the FACS labels (Methods), Cluster 2, Cluster 5, and Cluster 6 correspond to the common myeloid progenitors (CMP), megakaryocyte-erythrocyte progenitors (MEP) and granulocyte-monocyte progenitors (GMP), respectively (**Fig. 4c** and **Fig. 5a-c**). Cluster 1 is composed of long-term hematopoietic stem and progenitor cells (LTHSC) and multi-potent progenitors (MPP). These are cells from the early stage of differentiation. Cluster 4 consists of a mixture of MEP and CMP, while Cluster 3 is dominated by cells labeled as “other”. Comparison between the subpanel for BUSseq in **Fig. 4c** and **Fig. 5b** indicates that Cluster 4 are cells from an intermediate cell type between CMP and MEP. In particular, according to **Fig. 5e**, the marker genes *Apoe* and *Gata2* are highly expressed in Cluster 4 but not in CMP (Cluster 2) and MEP (Cluster 6), and the marker gene *Ctse* is expressed in MEP (Cluster 6) but not in Cluster 4 and CMP (Cluster 2). Therefore, cells in Cluster 4 do form a unique group with distinct expression patterns. This intermediate cell stage between CMP and GMP is missed by all of the other methods considered. Moreover, we find that well known B-cell lineage genes [36], *Ebf1, Vpreb1, Vpreb3*, and *Igll1*, are highly expressed in Cluster 3, but not in the other clusters (**Fig. 5c and e**). To identify Cluster 3, which is dominated by cells labeled as “other” by Nestorowa et al. [34], we map the mean expression profile of each cluster learned by BUSseq to the Haemopedia RNA-seq dataset [37]. It turns out that Cluster 3 aligns well to common lymphoid progenitors (CLP) that give rise to T-lineage cells, B-lineage cells and natural killer cells (**Fig. 5d**). Therefore, Cluster 3 represents cells that differentiate from lymphoid-primed multipotent progenitors (LMPP) cells [35]. Once again, all the other methods fail to identify these cells as a separate group.

**Figure 5:**
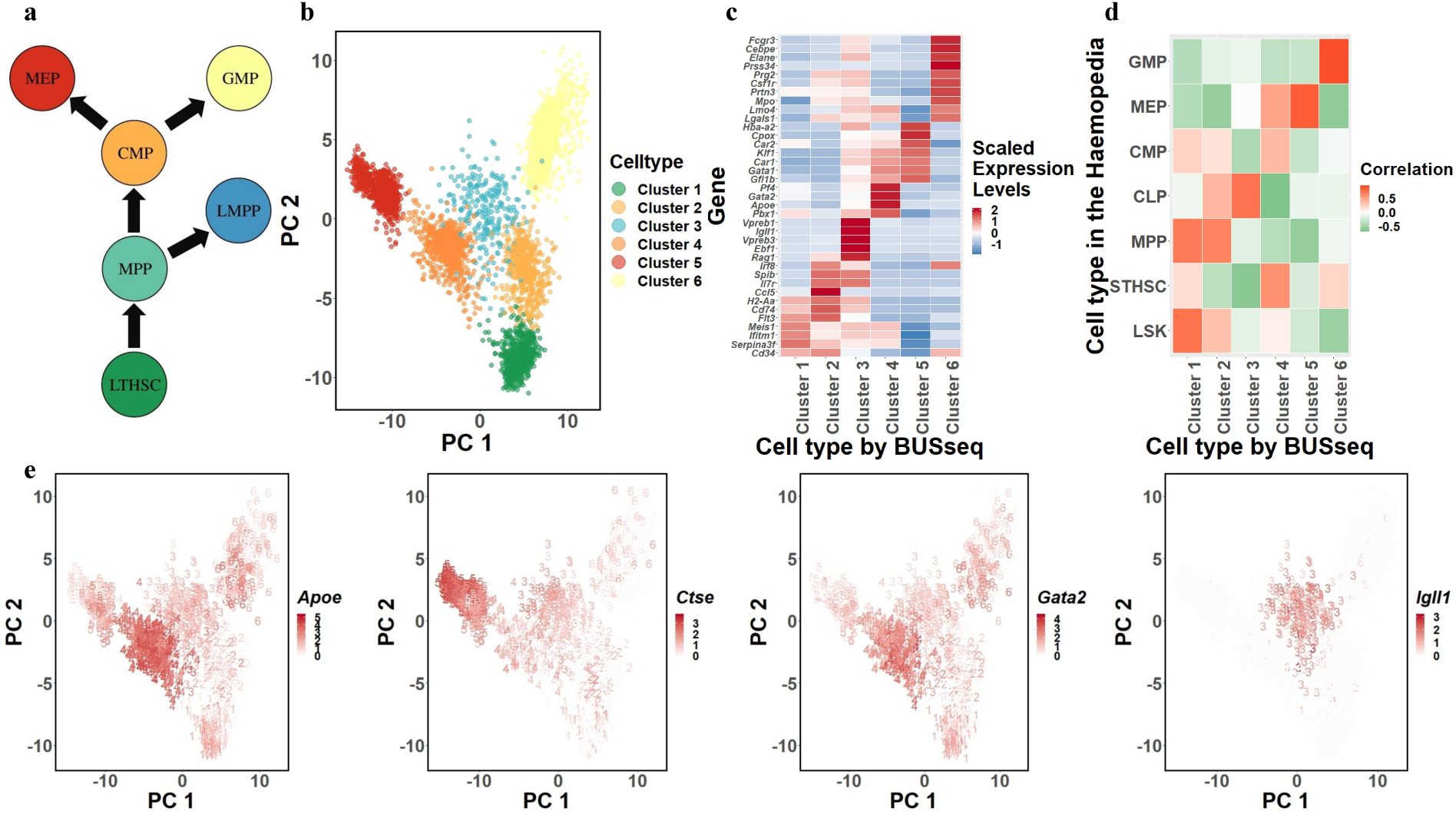
BUSseq preserves the hematopoietic stem and progenitor cells (HSPC) differentiation trajectories. **(a)** The diagram of HSPC differentiation trajectories. **(b)** The PCA plot of the corrected count matrix from BUSseq colored according to the estimated cell types by BUSseq. **(c)** The heatmap of scaled expression levels of key genes for HSPC. **(d)** The heatmap of correlation between gene expression profiles of each cell type inferred by BUSseq and those in the Haemopedia RNA-seq datasets. **(e)** The expression levels of four marker genes, *Apoe, Gata2, Ctse* and *Igll1*, shown in the PCA plots of corrected count data by BUSseq, respectively. The digit labels denote the corresponding clusters identified by BUSseq.

Thus, although BUSseq does not assume any temporal ordering between cell types, it is able to preserve the differentiation trajectories (**Fig. 5a and b**); although BUSseq assumes that each cell belongs to one cell type rather than conducts semisoft clustering [38], it is capable of capturing the subtle changes across cell types and within a cell type due to continuous processes such as development and differentiation.

We further inspect the functions of the intrinsic genes that distinguish different cell types. BUSseq detects 1419 intrinsic genes at the Bayesian FDR cutoff of 0.05 (Methods). The gene set enrichment analysis [39] shows that 51 KEGG pathways [40] are enriched among the intrinsic genes (p-values < 0.05) (Supplementary Notes). The highest ranked pathway is the Hematopoietic Cell Lineage Pathway, which corresponds to the exact biological process studied in the two datasets. Among the remaining 50 pathways, thirteen are related to the immune system, and another nine are associated with cell growth and differentiation (**Supplementary Table 1**). Therefore, the pathway analysis demonstrates that BUSseq is able to capture the underlying true biological variability, even if the batch effects are severe, as shown in **Fig. 3a** and **Fig. 4a**.

### BUSseq outperforms existing method on pancreas data

We further studied the four scRNA-seq datasets of human pancreas cells analyzed in Haghverdi et al. [13], two profiled by CEL-seq2 protocol [41, 42] and two assayed by SMART-seq2 protocol [42, 43]. These cells were isolated from deceased organ donors with and without type 2 diabetes. We obtained 7,095 cells after quality control (Supplementary Notes) and treated each dataset as a batch following Haghverdi et al. [13].

For the two datasets profiled by the SMART-seq2 protocol, Segerstolpe et al. [43] and Lawlor et al. [42] provide cell-type labels; for the other two datasets assayed by the CEL-seq2 protocol, Haghverdi et al. [13] provide the cell-type labels based on the marker genes in the original publications [41, 42]. We can thus compare the clustering results from each batch effects correction method with the labeled cell types (**Fig. 6a and b**).

**Figure 6:**
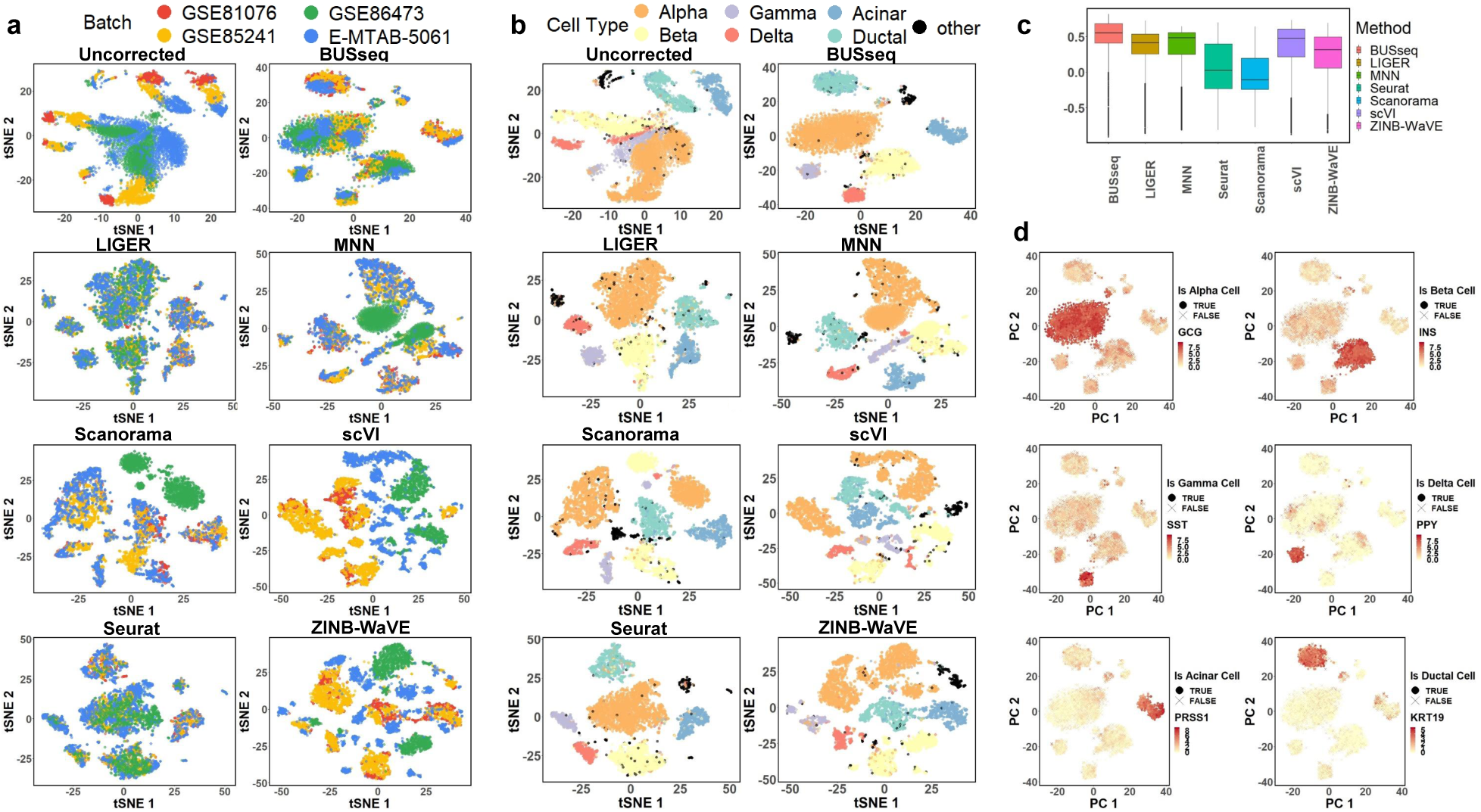
t-SNE plots for the pancreas data. **(a)** t-SNE plots colored by batch. **(b)** t-SNE plot colored by FACS cell type labels. **(c)** The boxplot of silhouette coefficients for all of the compared methods. **(d)** The expression levels of six marker genes, *GCG* for alpha cells, *INS* for beta cells, *SST* for gamma cells, *PPY* for delta cells, *PRSS1* for acinar cells, and *KRT19* for ductal cells, shown in the t-SNE plot of the corrected count data of BUSseq, respectively.

The pancreas is highly heterogeneous and consists of two major categories of cells: islet cells and non-islet cells. Islet cells include alpha, beta, gamma, and delta cells, while non-islet cells include acinar and ductal cells. BUSseq identifies a total of eight cell types: five for islet cells, two for non-islet cells and one for the labeled “other” cells. Specifically, the five islet cell types identified by BUSseq correspond to three groups of alpha cells, a group of beta cells, and a group of delta and gamma cells. The two non-islet cell types identified by BUSseq correspond exactly to the acinar and ductal cells. Compared to all of the other methods, BUSseq gives the best separation between islet and non-islet cells, as well as the best segregation within islet cells. In particular, the median silhouette coefficient by BUSseq is higher than that of any other method (**Fig. 6c**).

The ARIs of all methods are 0.608 for BUSseq, 0.542 for LIGER, 0.279 for MNN, 0.527 for Scanorama, 0.282 for scVI, 0.287 for Seurat and 0.380 for ZINB-WaVE. Thus, BUSseq outperforms all of the other methods in being consistent with the cell-type labels according to marker genes. In **Fig. 6d**, the locally high expression levels of marker genes for each cell type show that BUSseq correctly clusters cells according to their biological cell types.

BUSseq identifies 426 intrinsic genes at the Bayesian FDR cutoff of 0.05 (Methods). We conducted the gene set enrichment analysis [39] with the KEGG pathways [40] (Supplementary Notes). There are 14 enriched pathways (p-values < 0.05). Among them, three are diabetes pathways; two are pancreatic and insulin secretion pathways; and another two pathways are related to metabolism (**Supplementary Table 2**). Recall that the four datasets assayed pancreas cells from type 2 diabetes and healthy individuals, therefore, the pathway analysis once again confirms that BUSseq provides biologically and clinically valid cell typing.

## Discussion

For the completely randomized experimental design, it seems that “everyone is talking, but no one is listening.” Due to time and budget constraints, it is always difficult to implement a completely randomized design in practice. Consequently, researchers often pretend to be blind to the issue when carrying out their scRNA-seq experiments. In this paper, we mathematically prove and empirically show that under the more realistic reference panel and chain-type designs, batch effects can also be adjusted for scRNA-seq experiments. We hope that our results will alarm researchers of confounded experimental designs and encourage them to implement valid designs for scRNA-seq experiments in real applications.

BUSseq provides one-stop services. In contrast, most existing methods are multi-stage approaches—clustering can only be performed after the batch effects have been corrected and the differential expressed genes can only be called after the cells have been clustered. The major issue with multi-stage methods is that uncertainties in the previous stages are often ignored. For instance, when cells have been first clustered into different cell types and then differential gene expression identification is conducted, the clustering results are taken as if they were the underlying truth. As the clustering results may be prone to errors in practice, this can lead to false positives and false negatives. In contrast, BUSseq simultaneously corrects batch effects, clusters cell types, imputes missing data, and identifies intrinsic genes that differentiate cell types. BUSseq thus accounts for all uncertainties and fully exploits the information embedded in the data. As a result, BUSseq is able to capture subtler changes between cell types, such as the cluster corresponding to LMPP lineage that is missed by all the state-of-the-art methods.

BUSseq employs MCMC for statistical inference. As MCMC algorithms not only provide point estimates but also explore the entire posterior distributions and hence allow the users to quantify the uncertainty of estimates, they are famous for heavy computation load. However, fortunately, the computational complexity of BUSseq is 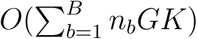, which is both linear in the number of genes *G* and in the total number of cells 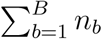. Moreover, most steps of the MCMC algorithm for BUSseq are parallelizable. We implement a parallel multi-core-CPU version and a parallel GPU version of the algorithm, respectively. Running the GPU version of the algorithm with a single core of an Intel Xeon Gold 6132 Processor and one NVIDIA Tesla P100 GPU took 0.35, 1.15, 1.5 hours for the simulation, the hematopoietic and the human pancreas data, respectively (**Supplementary Table 3**). Compared to the time for preparing samples and conducting the scRNA-seq experiments, the computation time of BUSseq is affordable and worthwhile for the accuracy.

Practical and valid experimental designs are urgently required for scRNA-seq experiments. We envision that the flexible reference panel and the chain-type designs will be widely adopted in scRNA-seq experiments and BUSseq will greatly facilitate the analysis of scRNA-seq data.

## Acknowledgments

This work was supported by the Hong Kong Ph.D. Fellowship PF15-17417 and the General Research Funds 14306417 and 14305319 from the Hong Kong Research Grants Council of the Hong Kong Special Administrative Region of the People’s Republic of China and Direct Grants from the Research Committee of the Chinese University of Hong Kong. We acknowledge Dr. Xiangyu Luo for helpful comments on an early version of our paper.

## Author Contributions

FD.S developed the method and the proof, implemented the algorithm, prepared the software package, analyzed the data, and wrote the paper. GM.C. implemented the algorithm and analyzed the data. YY.W. conceived and supervised the study, developed the method and the proof, and wrote the paper.

## Methods

### BUSseq model

The hierarchical model of BUSseq can be summarized as:

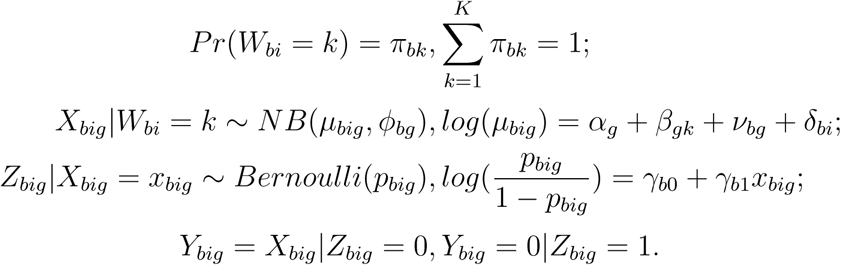

Collectively, 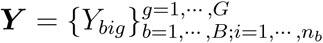 are the observed data; the underlying expression levels 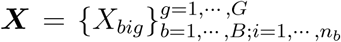, the dropout indicators 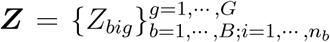 and the cell type indicators 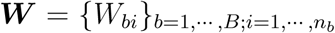 are all missing data; the log-scale baseline gene expression levels ***α*** = *{α*_*g*_}_*g*=1,…, *G*_, the cell type effects 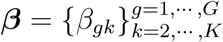, the location batch effects 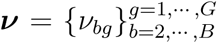, the overdispersion parameters 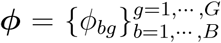, the cell-specific size factors 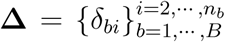, the dropout parameters **Γ** = {*γ*_*b*0_, *γ*_*b*1_}^*b*=1, …, *B*^ and the cell compositions 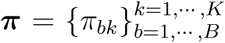 are the parameters. Without loss of generality, for model identifiability, we assume that the first batch is the reference batch measured without batch effects with 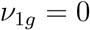 for every gene and the first cell type is the baseline cell type with *β*_*g*1_ = 0 for every gene. Similarly, we take the cell-specific size factor *δ*_*b*1_ = 0 for the first cell of each batch. We gather all the parameters as **Θ** = {***α, β, ν, ϕ*, Δ, Γ, *π***}.

### Experimental designs

By creating a set of functions similar to the probability generating function, we prove that BUSseq is identifiable, in other words, if two sets of parameters are different, then their probability distribution functions for the observed data are different, for not only the “complete setting” but also the “reference panel” and the “chain-type” designs (see the proofs in the Supplementary Notes).

#### Theorem 1. (The Complete Setting)

*If π*_*bk*_ > 0 *for every batch b and cell type k, given that (I) γ*_*b*1_ < 0 *for every b, (II) for any two cell types k*_1_ *and k*_2_, *there exist at least two differentially expressed genes g*_1_ *and g*_2_*—* 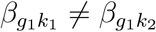 *and* 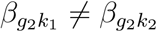, *and (III) for any two distinct cell-type pairs* (*k*_1_, *k*_2_) ≠ (*k*_3_, *k*_4_), *their differences in cell-type effects are not the same* 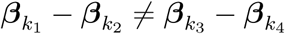, *then BUSseq is identifiable (up to label switching) in the sense that L*_*o*_(**Θ**|***y***) = *L*_*o*_(**Θ**^*\**^|***y***) *for any* ***y*** *implies that* 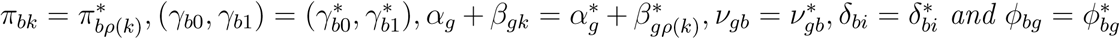 *for every gene g and batch b, where ρ is a permutation of {*1, 2, …, *K}.*

In the following, we denote the cell types that are present in batch *b* as *C*_*b*_ and count the number of cell types existing in batch *b* as *K*_*b*_ = |*C*_*b*_|.

#### Theorem 2. (The Reference Panel Design)

*If there are a total of K cell types* 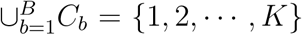 *K*_*b*_ ≥ 2 *for every batch b, and there exists a batch* 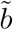 *such that it contains all of the cell types* 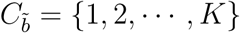, *then given that conditions (I)-(III) hold, BUSseq is identifiable (up to label switching).*

#### Theorem 3. (The Chain-type Design)

*If there are a total of K cell types* 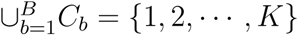 *and every two consecutive batches share at least two cell types* |*C*_*b*_ *∩ C*_*b*−1_| *≥* 2 *for all b ≥* 2, *then given that conditions (I)-(III) hold, BUSseq is identifiable (up to label switching).*

Therefore, even for the “reference panel” and “chain-type” designs that do not assay all cell types in each batch, batch effects can be removed; cell types can be clustered; and missing data due to dropout events can be imputed. Both the reference panel design and the chain-type design belong to the more general connected design.

#### Theorem 4. (The Connected Design)

*We define a batch graph G* = (*V, E*). *Each node b* ∈ *V represents a batch. There is an edge e* ∈ *E between two nodes b*_1_ *and b*_2_ *if and only if batches b*_1_ *and b*_2_ *share at least two cell types. If the batch graph is connected and conditions (I)-(III) hold, then BUSseq is identifiable (up to label switching).*

### Statistical inference

We conduct the statistical inference under the Bayesian framework. We assign independent priors to each component of **Θ** as follows: 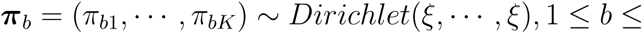 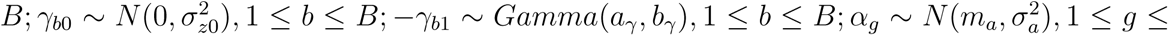 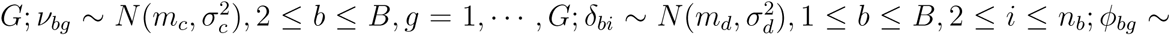 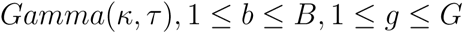.

We are interested in detecting genes that differentiate cell types. Therefore, we impose a spike-and-slab prior [44] using a normal mixture to the cell-type effect *β*_*gk*_. The spike component concentrates on zero with a small variance 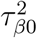, whereas the slab component tends to deviate from zero, thus having a larger variance 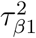. We introduce another latent variable *L*_*gk*_ to indicate which component *β*_*gk*_ comes from. *L*_*gk*_ = 0 if gene *g* is not differentially expressed between cell type *k* and cell type one, and *L*_*gk*_ = 1, otherwise. We further define 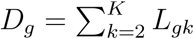. If *D*_*g*_ > 0, then the expression level of gene *g* does not stay the same across cell types. Following Huo et al. [12], we call such genes intrinsic genes, which differentiate cell types. To control for multiple hypothesis testing, we let *L*_*gk*_ ~ *Bernoulli*(*p*) and assign a conjugate prior *Beta*(*a*_*p*_, *b*_*p*_) to *p*. We set *τ*_*β*1_ to a large number and let *τ*_*β*0_ follow an inverse-gamma prior *Inv* − *Gamma*(*a*_*τ*_, *b*_*τ*_) with a small prior mean.

We develop an MCMC algorithm to sample from the posterior distribution (Supplementary Notes). After the burn-in period, we take the mean of the posterior samples to estimate ***γ***_*b*_, *α*_*g*_, *β*_*gk*_, *ν*_*bg*_, *δ*_*bi*_ and *ϕ*_*bg*_ and use the mode of posterior samples of *W*_*bi*_ to infer the cell type for each cell.

When inferring the differential expression indicator *L*_*gk*_, we control the Bayesian false discovery rate (FDR) [32] defined as

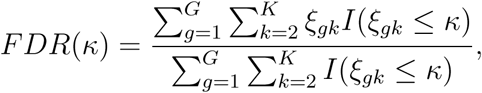

where *ξ* _*gk*_ = *Pr*(*L*_*gk*_ = 0|***y***) is the posterior marginal probability that gene *g* is not differentially expressed between cell type *k* and cell type one, which can be estimated by the *T* posterior samples 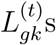 s collected after the burn-in period as 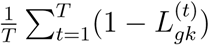). Given a control level *α* such as 0.1, we search for the largest *κ*_0_ *≤* 0.5 such that the estimated 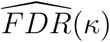 based on 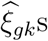 is smaller than *α* and declare 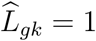 if 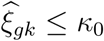. The upper bound 0.5 for *κ*_0_ prevents us from calling differentially expressed genes with small posterior probability *Pr*(*L*_*gk*_ = 1|***y***). Consequently, we identify the genes with 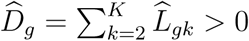 as the intrinsic genes. We set *α* = 0.05 in both the simulation study and the real applications.

BUSseq allows the user to input the total number of cell types *K* according to prior knowledge. When *K* is unknown, BUSseq selects the number of cell types 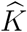 such that it achieves the minimum BIC (Supplementary Notes).

### Batch-effects-corrected values

To facilitate further downstream analysis, we also provide a version of count data 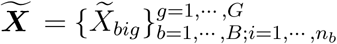 for which the batch effects are removed and the biological variability is retained. We develop a quantile matching approach based on inverse sampling. Specifically, given the fitted model and the inferred underlying expression level 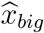, we first sample *u*_*big*_ from 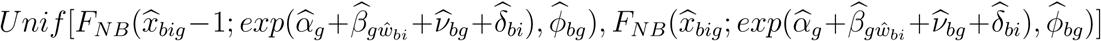 where *Unif* [*a, b*] denotes the uniform distribution on the interval [*a, b*] and *F*_*NB*_(·; *µ, r*) denotes the cumulative distribution function of a negative binomial distribution with mean *µ* and overdispersion parameter *r*. Next, we calculate the 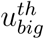 quantile of 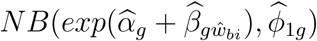 as the corrected value 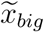.

The corrected data 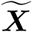 are not only protected from batch effects but also impute the missing data due to dropout events. Moreover, further cell-specific normalization is not needed. Meanwhile, the biological variability is retained thanks to the quantile transformation and sampling step. Therefore, we can directly perform downstream analysis on 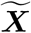.

### Assignment of FACS cell type labels to learned clusters

In the two real data examples, we first identify the cell type of each individual cell according to FACS labeling. Then, for each cluster learned by BUSseq, we calculate the proportion of labeled cell types. If a cell type accounts for more than one-third of the cells in a given cluster, we assign this cell type to the cluster. Although a cluster may be assigned more than one cell type, most identified clusters by BUSseq are dominated by only one cell type.

### Mapping clusters to Haemopedia

Haemopedia is a database of gene expression profiles from diverse types of hematopoietic cells [37]. It collected flow sorted cell populations from healthy mice.

To understand Cluster 3 learned by BUSseq for the hematopoietic data, which is dominated by cells classified as “other” according to the FACS labeling, we mapped the cluster means learned by BUSseq to the Haemopedia RNA-seq dataset.

We first applied TMM normalization [45] to all the samples in the Haemopedia RNA-seq dataset. Then, we extracted 7 types of hematopoietic stem and progenitor cells from Haemopedia, including Lin^−^Sca-1^+^c-Kit^+^ (LSK) cells, short-term hematopoietic stem cells (STHSC), MPP, CLP, CMP, MEP and GMP. Each selected cell type had two RNA-seq samples in Haemopedia, so we averaged over the two replicates for each cell type. Further, we added one to the normalized expression levels as a pseudo read count to handle genes with zero read count and log-transformed the data. Finally, we scaled the data across the 7 cell types for each gene. To be comparable, we transformed the cluster mean learned by BUSseq as *m*_*gk*_ = *log*(1 + *exp*(*α*_*g*_ + *β*_*gk*_)) for gene *g* in the cluster *k* and scaled *m*_*gk*_ across all cell types as well. Finally, we calculated the correlation between the cluster means inferred by BUSseq and the reference expression profiles in Haemopedia for 37 marker genes. The 37 marker genes were retrieved from Paul et al. [35] (31 maker genes for HSPC) and Herman et al. [36] (6 maker genes for LMPP).

### Software availability

The C++ source code of the parallel multi-core-CPU version of BUSseq is available on GitHub https://github.com/songfd2018/BUSseq-1.0, and the CUDA C source code of the GPU version of BUSseq is available on GitHub https://github.com/Anguscgm/BUSseq_gpu. All codes for producing results and figures in this manuscript are also available on GitHub (https://github.com/songfd2018/BUSseq-1.0_implementation).

### Data availability

The published data sets used in this manuscript are available through the following accession numbers: SMART-seq2 platform hematopoietic data with GEO GSE81682 by Nestorowa et al. [34]; MARS-seq platform hematopoietic data with GEO GSE72857 by Paul et al. [35]; CEL-seq platform pancreas data with GEO GSE81076 by Grün et al. [41]; CEL-seq2 platform pancreas data with GEO GSE85241 by Muraro et al. [46]; SMART-seq2 platform pancreas data with GEO GSE86473 by Lawlor et al. [42]; and SMART-seq2 platform pancreas data with ArrayExpress E-MTAB-5061 by Segerstolpe et al. [43].

The parameter settings for the simulation study and the simulated data are available on GitHub (https://github.com/songfd2018/BUSseq-1.0_implementation).

