## Supplementary Materials for "Flexible Experimental Designs for Valid Single-cell RNA-sequencing Experiments Allowing Batch Effects Correction"

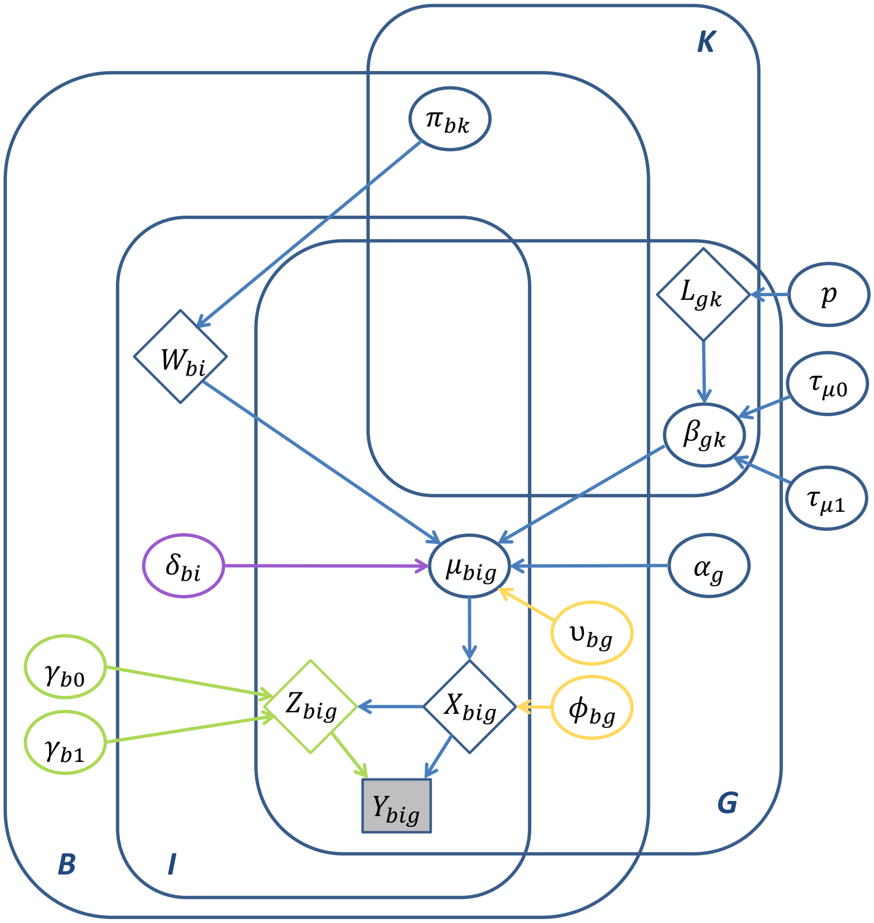


Supplementary Figure 1: The graphical representation of the BUSseq model. The yellow color corresponds to batch effects; the green color models the dropout events; the purple color indicates the cell-specific size factor. The ellipses are for parameters; the diamonds represent latent variables; and only $Y_{big}$ in the grey rectangle is observed.

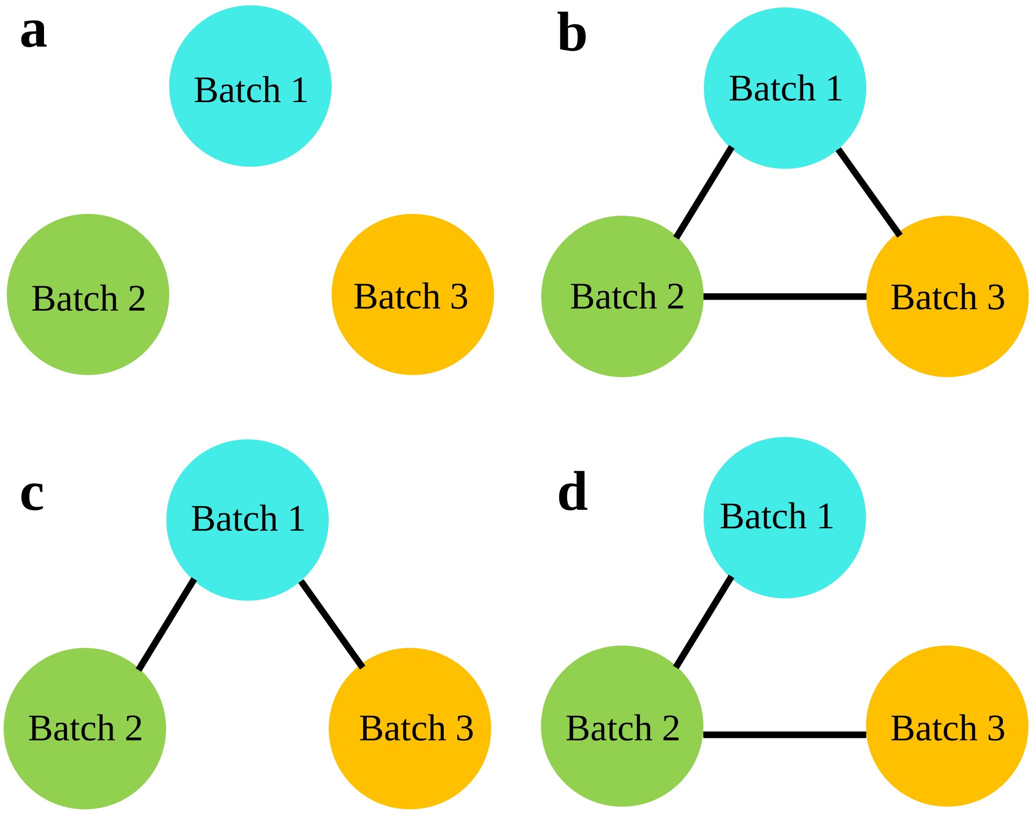


Supplementary Figure 2: The batch graphs for the experiment designs in **Figure 2**. **(a)** The confounded design. **(b)** The complete setting design. **(c)** The reference panel design. **(d)** The chain-type design.

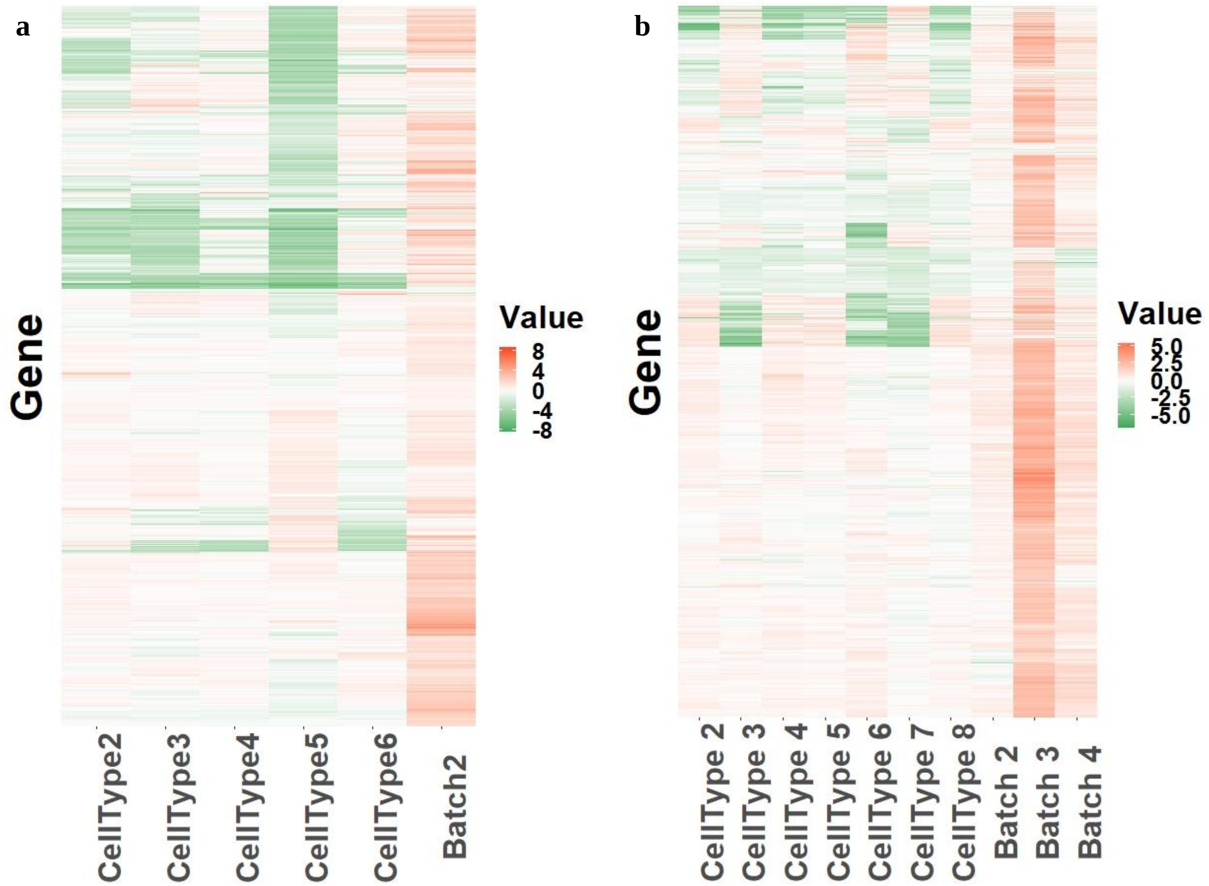


Supplementary Figure 3: The heatmap of the estimated cell type effects and batch effects by BUSseq in two real studies. Each row represents a gene, and each column corresponds to a kind of the cell type effect $\beta_{gk}, 2 \leq k \leq K$ or the batch effect $\nu_{bg}, 2 \leq b \leq B$. **(a)** Mouse Hematopoietic study. **(a)** Human Pancreas study.

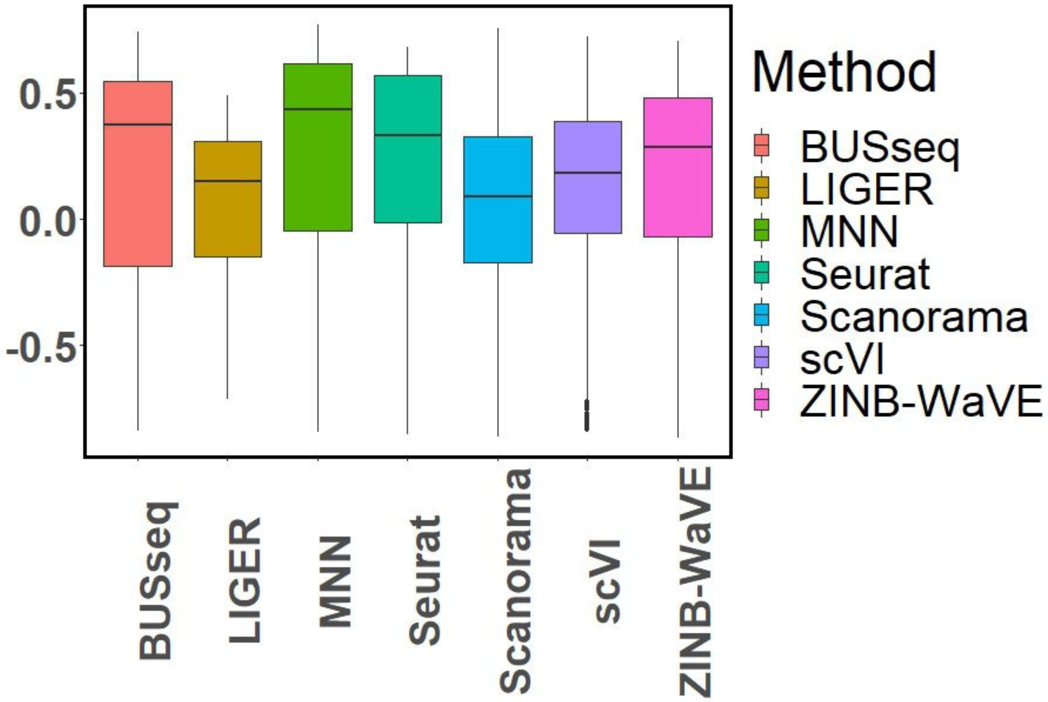


Supplementary Figure 4: The boxplots of silhouette coefficients for all of the compared methods in the hematopoietic study.

| Ranking | Pathway | p values | Category |
|---|---|---|---|
| 1 | Hematopoietic cell lineage | $1.73 \times 10^{-14}$ | |
| 2 | Cytokine-cytokine receptor interaction | $1.84 \times 10^{-12}$ | Cell growth and differentiation |
| 3 | Cell adhesion molecules (CAMs) | $3.29 \times 10^{-9}$ | Immune system |
| 4 | Leukocyte transendothelial migration | $1.54 \times 10^{-6}$ | Immune system |
| 5 | Primary immunodeficiency | $6.75 \times 10^{-6}$ | Immune system |
| 6 | Rap1 signaling pathway | $3.44 \times 10^{-5}$ | Cell growth and differentiation |
| 7 | Transcriptional misregulation in cancer | $4.23 \times 10^{-5}$ | |
| 8 | Rheumatoid arthritis | $6.59 \times 10^{-5}$ | |
| 9 | Pathways in cancer | $1.15 \times 10^{-4}$ | |
| 10 | Tuberculosis | $1.40 \times 10^{-4}$ | |
| 11 | Malaria | $3.02 \times 10^{-4}$ | |
| 12 | Toll-like receptor signaling pathway | $3.60 \times 10^{-4}$ | |
| 13 | Staphylococcus aureus infection | $4.53 \times 10^{-4}$ | |
| 14 | PI3K-Akt signaling pathway | $5.53 \times 10^{-4}$ | Cell growth and differentiation |
| 15 | Osteoclast differentiation | $9.74 \times 10^{-4}$ | Cell growth and differentiation |
| 16 | T cell receptor signaling pathway | $9.96 \times 10^{-4}$ | Immune system |
| 17 | Intestinal immune network for IgA production | $1.43 \times 10^{-3}$ | Immune system |
| 18 | Leishmaniasis | $1.44 \times 10^{-3}$ | Immune system |
| 19 | Platelet activation | $1.62 \times 10^{-3}$ | |
| 20 | NF-kappa B signaling pathway | $1.65 \times 10^{-3}$ | Immune system |
| 21 | Asthma | $2.13 \times 10^{-3}$ | |
| 22 | Jak-STAT signaling pathway | $2.61 \times 10^{-3}$ | Cell growth and differentiation |
| 23 | B cell receptor signaling pathway | $3.34 \times 10^{-3}$ | Immune system |
| 24 | ECM-receptor interaction | $3.99 \times 10^{-3}$ | Cell growth and differentiation |
| 25 | Neuroactive ligand-receptor interaction | $4.05 \times 10^{-3}$ | |
| 26 | Ras signaling pathway | $4.93 \times 10^{-3}$ | Cell growth and differentiation |
| 27 | Pertussis | $5.48 \times 10^{-3}$ | |
| 28 | Inflammatory bowel disease (IBD) | $6.51 \times 10^{-3}$ | Immune system |
| 29 | Thyroid hormone signaling pathway | $9.06 \times 10^{-3}$ | |
| 30 | Phagosome | $9.51 \times 10^{-3}$ | Immune system |
| 31 | Mineral absorption | $1.09 \times 10^{-2}$ | |
| 32 | Amoebiasis | $1.17 \times 10^{-2}$ | |
| 33 | Focal adhesion | $1.43 \times 10^{-2}$ | Cell growth and differentiation |
| 34 | Glycosphingolipid biosynthesis - lacto and neolacto series | $1.49 \times 10^{-2}$ | |
| 35 | p53 signaling pathway | $1.67 \times 10^{-2}$ | |
| 36 | Calcium signaling pathway | $1.70 \times 10^{-2}$ | |
| 37 | Fc epsilon RI signaling pathway | $1.86 \times 10^{-2}$ | |
| 38 | Natural killer cell mediated cytotoxicity | $1.98 \times 10^{-2}$ | Immune system |
| 39 | Proteoglycans in cancer | $2.02 \times 10^{-2}$ | |
| 40 | Chemokine signaling pathway | $2.40 \times 10^{-2}$ | Immune system |
| 41 | Gastric acid secretion | $2.74 \times 10^{-2}$ | |
| 42 | ABC transporters | $2.82 \times 10^{-2}$ | |
| 43 | HIF-1 signaling pathway | $3.25 \times 10^{-2}$ | |
| 44 | Chagas disease (American trypanosomiasis) | $3.50 \times 10^{-2}$ | |
| 45 | Retrograde endocannabinoid signaling | $3.50 \times 10^{-2}$ | |
| 46 | NOD-like receptor signaling pathway | $3.63 \times 10^{-2}$ | Immune system |
| 47 | Aldosterone-regulated sodium reabsorption | $3.77 \times 10^{-2}$ | |
| 48 | Sphingolipid signaling pathway | $3.87 \times 10^{-2}$ | |
| 49 | Progesterone-mediated oocyte maturation | $4.41 \times 10^{-2}$ | |
| 50 | MAPK signaling pathway | $4.58 \times 10^{-2}$ | Cell growth and differentiation |
| 51 | Carbohydrate digestion and absorption | $4.76 \times 10^{-2}$ | |

Supplementary Table 1: The 51 KEGG pathways (p-value $< 0.05$ [1]) significantly enriched among the intrinsic genes identified by BUSseq from the hematopoietic data.

| Ranking | Pathway | p values | |
|---|---|---|---|
| 1 | Maturity onset diabetes of the young | $9.09 \times 10^{-9}$ | Diabetes |
| 2 | Pancreatic secretion | $6.42 \times 10^{-7}$ | Protein Secretion |
| 3 | Insulin secretion | $1.58 \times 10^{-6}$ | Protein Secretion |
| 4 | Protein digestion and absorption | $1.89 \times 10^{-3}$ | Metabolism |
| 5 | ECM-receptor interaction | $7.08 \times 10^{-3}$ | |
| 6 | Type II diabetes mellitus | $7.63 \times 10^{-3}$ | Diabetes |
| 7 | Morphine addiction | $8.99 \times 10^{-3}$ | |
| 8 | Proteoglycans in cancer | $1.44 \times 10^{-2}$ | |
| 9 | Dopaminergic synapse | $1.76 \times 10^{-2}$ | |
| 10 | GABAergic synapse | $2.24 \times 10^{-2}$ | |
| 11 | Type I diabetes mellitus | $2.26 \times 10^{-2}$ | Diabetes |
| 12 | Tight junction | $2.48 \times 10^{-2}$ | |
| 13 | Drug metabolism - cytochrome P450 | $3.08 \times 10^{-2}$ | Metabolism |
| 14 | Focal adhesion | $4.07 \times 10^{-2}$ | |

Supplementary Table 2: The 14 KEGG pathways (p-value$<$ 0.05 [1]) significantly enriched among the intrinsic genes identified by BUSseq from the pancreatic data.

| Study | Simulation | Mouse Hematpoietic | Human Pancreas |
|---|---|---|---|
| Number of cell $N$ | 1,000 | 4,649 | 7,095 |
| Number of highly variable genes $G$ | 3,000 | 3,470 | 2,480 |
| Number of batches $B$ | 4 | 2 | 4 |
| Number of cell types $K$ | 5 | 6 | 8 |
| Total number of iterations in MCMC | 4,000 | 8,000 | 8,000 |
| Number of burn-in iterations | 2,000 | 4,000 | 4,000 |
| CPU Runtime (hours) | 1.00 | 8.00 | 12.09 |
| GPU Runtime (hours) | 0.35 | 1.15 | 1.50 |

Supplementary Table 3: The summary and runtime of BUSseq in the simulation and case studies. The CPU runtime records the time consumption of running BUSseq on eight Intel Xeon Gold 6128 CPU cores in parallel, whereas the GPU runtime records the time consumption of running BUSseq with a single core of an Intel Xeon Gold 6132 CPU of 512GM RAM and one NVIDIA Tesla P100 GPU of 16GM RAM.

### Supplementary Notes

#### Processing of the real datasets

For the two hematopoietic datasets, we downloaded the read count matrix of the 1,920 cells profiled by Paul et al. [2] and the 2,729 cells labeled as myeloid progenitor cells by Nestorowa et al. [3] from the NCBI Gene Expression Omnibus (GEO) with the accession numbers GSE72857 and GSE81682. Following Brennecke et al. [4], we sorted the genes according to their adjusted variance-mean ratio of expression levels in both datasets separately and focused on the 3,470 genes that are highly variable in both datasets.

Two of the pancreas datasets profiled by the CEL-seq2 platform were downloaded from GEO with accession number GSE80176 [5] and GSE86473 [6]. The two datasets assayed by the SMART-seq2 platform were obtained from GSE85241 [7] and from ArrayExpress accession number E-MATB-5061 [8]. Following Haghverdi et al. [9], we excluded cells with low library sizes ($< 100,000$ reads), low numbers of expressed genes ($> 40\%$ total counts from ribosomal RNA genes), or high ERCC content ($> 20\%$ of total counts from spike-in transcripts) resulting in 7,095 cells. We selected the 2,480 highly variable genes shared by the four datasets according to Brennecke et al. [4] by sorting the ratio of variance and mean expression level after adjusting technical noise with the variances of spike-in transcripts. The cell types of the two datasets profiled by the CEL-seq2 platform were labeled according to Lawlor et al. [6] and Grün et al. [5], with the GCG gene marking alpha islets, INS for beta islets, SST for delta islets, PPY for gamma islets, PRSS1 for acinar cells, and KRT19 for ductal cells. The cell types of the other two datasets assayed by the SMART-seq2 platform were provided in their metadata.

#### Comparison measures

The adjusted Rand index (ARI) measures the consistency between two clustering results and is between zero and one, a higher value indicating better consistency. It is defined as:

$$ARI = \frac{\sum_{i=1}^{I}\sum_{j=1}^{J}\binom{n_{ij}}{2} - [\sum_{i=1}^{I}\binom{a_i}{2}\cdot\sum_{j=1}^{J}\binom{b_j}{2}]\binom{n}{2}}{(1/2)[\sum_{i=1}^{I}\binom{a_i}{2} + \sum_{j=1}^{J}\binom{b_j}{2}] - [\sum_{i=1}^{I}\binom{a_i}{2}\cdot\sum_{j=1}^{J}\binom{b_j}{2}]\binom{n}{2}},$$

where $n_{ij}, a_i, b_j$ are values from the contingency table of two clusterings. ARIs are calculated by R package *mclust* [10].

To evaluate the separation of different cell types after correction, we calculate the silhouette coefficient of each cell using the R package *cluster* [11]. We regard each cell type, either the truth known in the simulation study or the labeling according to FACS, as a cluster. Let $a(i)$ be the average distance of cell $i$ to all the other cells assigned to the same cluster as cell $i$, and let $b(i)$ be the average distance of cell $i$ to all cells in the neighboring cluster, i.e., the cluster with the lowest average distance to cell $i$'s cluster. The silhouette coefficient for cell $i$ is defined as:

$$s(i) = \frac{b(i) - a(i)}{min(a(i), b(i))}.$$

The silhouette coefficient $s(i)$ ranges from -1 to 1. The larger the values of $s(i)$, the closer cell $i$ is to cells in the same cluster than cells in other clusters. We calculate the silhouette coefficient according to the t-SNE coordinates obtained from the corrected count data matrix (BUSseq and MNN) or from low-dimensional representations (LIGER, Scanorama, scVI, Seurat and ZINBWaVE).

##### The benchmarked methods

To ensure a fair comparison, we follow the preprocessing steps of each of the methods used for benchmarking according to their original publications. LIGER [12] normalizes the raw read count of each cell by the cell's total read counts (see `https://github.com/MacoskoLab/liger`). MNN [9] takes the first batch as the reference batch and normalizes the other batches to adjust for difference in sequencing depths (see `https://github.com/MarioniLab/MNN2017`). Scanorama [13] conducts $L_2$-normalization in the preprocessing steps (see `https://github.com/brianhie/scanorama`). Seurat [14] log-transforms and scales the observed read count data (see `https://satijalab.org/seurat/`). scVI [15] and ZINB-WaVE [16] directly work on the raw read count data (see `https://github.com/YosefLab/scVI` and `https://github.com/drisso/zinbwave`, respectively). For BUSseq, we run the MCMC algorithm for 4,000, 8,000 and 8,000 iterations for the simulated data, the hematopoietic study and the pancreas study, respectively. In each case, we treat the first half of all the iterations as the burn-in period and use the posterior samples collected from the second half for statistical inference. Please see `https://github.com/songfd2018/BUSseq-1.0_implementation` for the specification of hyperparameters used in this manuscript.

##### Implementation of pathway analysis

To identify the biological functions of the intrinsic genes, we conduct gene set enrichment analysis for the intrinsic genes on KEGG pathways using DAVID [1]. We control the Expression Analysis Systematic Explorer Score, a modified version of Fisher exact p-value, at the level of 0.05 to identify enriched pathways.

#### Details of the BUSseq model

##### BUSseq model

The hierarchical model of BUSseq can be summarized as:

$$\begin{aligned}
&Pr(W_{bi}=k)=\pi_{bk}, \sum_{k=1}^{K}\pi_{bk}=1;\\
&X_{big}|W_{bi}=k \sim NB(\mu_{big},\phi_{bg}), \quad log(\mu_{big})=\alpha_g+\beta_{gk}+\nu_{bg}+\delta_{bi};\\
&Z_{big}|X_{big}=x_{big} \sim Bernoulli(p_{big}), \quad log(\frac{p_{big}}{1-p_{big}})=\gamma_{b0}+\gamma_{b1}x_{big};\\
&Y_{big}=X_{big}|Z_{big}=0, \quad Y_{big}=0|Z_{big}=1. \qquad (1)
\end{aligned}$$

Collectively, $\boldsymbol{Y} = \{Y_{big}\}_{b=1,\cdots,B;i=1,\cdots,n_b}^{g=1,\cdots,G}$ are the observed data; the underlying expression levels $\boldsymbol{X} = \{X_{big}\}_{b=1,\cdots,B;i=1,\cdots,n_b}^{g=1,\cdots,G}$, the dropout indicators $\boldsymbol{Z} = \{Z_{big}\}_{b=1,\cdots,B;i=1,\cdots,n_b}^{g=1,\cdots,G}$ and the cell type indicators $\boldsymbol{W} = \{W_{bi}\}_{b=1,\cdots,B;i=1,\cdots,n_b}$ are all missing data; the log-scale baseline gene expression levels $\boldsymbol{\alpha} = \{\alpha_g\}_{g=1,\cdots,G}$, the cell type effects $\boldsymbol{\beta} = \{\beta_{gk}\}_{k=2,\cdots,K}^{g=1,\cdots,G}$, the location batch effects $\boldsymbol{\nu} = \{\nu_{bg}\}_{b=2,\cdots,B}^{g=1,\cdots,G}$, the overdispersion parameters $\boldsymbol{\phi} = \{\phi_{bg}\}_{b=1,\cdots,B}^{g=1,\cdots,G}$, the cell-specific size factors $\boldsymbol{\Delta} = \{\delta_{bi}\}_{b=1,\cdots,B}^{i=2,\cdots,n_b}$, the dropout parameters $\boldsymbol{\Gamma} = \{\gamma_{b0}, \gamma_{b1}\}^{b=1,\cdots,B}$ and the cell compositions $\boldsymbol{\pi} = \{\pi_{bk}\}_{b=1,\cdots,B}^{k=1,\cdots,K}$ are the parameters. Without loss of generality, for model identifiability, we assume that the first batch is the reference batch measured without batch effects with $\nu_{1g} = 0$ for every gene and the first cell type is the baseline cell type with $\beta_{g1} = 0$ for every gene. Similarly, we take the cell-specific size factor $\delta_{b1} = 0$ for the first cell of each batch. We gather all the parameters as $\boldsymbol{\Theta} = \{\boldsymbol{\alpha}, \boldsymbol{\beta}, \boldsymbol{\nu}, \boldsymbol{\phi}, \boldsymbol{\Delta}, \boldsymbol{\Gamma}, \boldsymbol{\pi}\}$. Let $f_{NB}(x; \mu, \phi) = C_x^{\phi+x-1}(\frac{\mu}{\mu+\phi})^x(\frac{\phi}{\mu+\phi})^\phi$ denote the probability mass function of the negative binomial distribution $NB(\mu, \phi)$, where $C_k^n$ is the binomial coefficient, then the complete data likelihood function equals to:

$$L_c(\boldsymbol{\Theta}|\boldsymbol{y}, \boldsymbol{x}, \boldsymbol{z}, \boldsymbol{w}) = \prod_{b=1}^{B}\prod_{i=1}^{n_b}\prod_{k=1}^{K}\{\pi_{bk}\prod_{g=1}^{G}[I(y_{big} = x_{big}(1 - z_{big}))\frac{exp[(\gamma_{b0} + \gamma_{b1}x_{big})z_{big}]}{1 + exp(\gamma_{b0} + \gamma_{b1}x_{big})}$$
$$\cdot f_{NB}(x_{big}; exp(\alpha_g + \beta_{gk} + \nu_{bg} + \delta_{bi}), \phi_{bg})]\}^{I(w_{bi}=k)}. \quad (2)$$

Consequently, the observed data likelihood function becomes

$$L_o(\boldsymbol{\Theta}|\boldsymbol{y}) = \prod_{b=1}^{B}\prod_{i=1}^{n_b}[\sum_{k=1}^{K}\pi_{bk}\prod_{g=1}^{G}Pr(Y_{big} = y_{big}|\boldsymbol{\Theta})], \quad (3)$$

$$Pr(Y_{big} = y_{big}|\boldsymbol{\Theta}) = \begin{cases} \sum_{x=1}^{\infty}\frac{exp(\gamma_{b0}+\gamma_{b1}x)}{1+exp(\gamma_{b0}+\gamma_{b1}x)}f_{NB}(x; exp(\alpha_g + \beta_{gk} + \nu_{bg} + \delta_{bi}), \phi_{bg}) & \\ +f_{NB}(0; exp(\alpha_g + \beta_{gk} + \nu_{bg} + \delta_{bi}), \phi_{bg}) & y_{big} = 0, \\ \frac{1}{1+exp(\gamma_{b0}+\gamma_{b1}y_{big})}f_{NB}(y_{big}; exp(\alpha_g + \beta_{gk} + \nu_{bg} + \delta_{bi}), \phi_{bg}) & y_{big} > 0. \end{cases}$$

#### The Markov chain Monte Carlo (MCMC) Algorithm for BUSseq

We develop an MCMC algorithm to sample from the posterior distribution (Supplementary Information). After the burn-in period, we take the mean of the posterior samples to estimate $\boldsymbol{\gamma}_b, \alpha_g, \beta_{gk}, \nu_{bg}, \delta_{bi}$ and $\phi_{bg}$ and use the mode of posterior samples of $W_{bi}$ to infer the cell type for each cell.

To conduct posterior inference, we develop an MCMC algorithm to draw samples from the posterior distribution. At iteration $t$:

1. Update $z_{big}^{[t]}$ and $x_{big}^{[t]}$ sequentially for $(b, i, g)$, if $y_{big} = 0$:

$$z_{big}^{[t]} \begin{cases} = 1 & , \text{if } x_{big}^{[t-1]} > 0; \\ \sim Bernoulli(\frac{exp(\gamma_{b0}^{[t-1]})}{1+exp(\gamma_{b0}^{[t-1]})}) & , \text{if } x_{big}^{[t-1]} = 0. \end{cases}$$

$$x_{big}^{[t]} \begin{cases} = 0 & , \text{if } z_{big}^{[t]} = 0; \\ \propto \frac{exp(\gamma_{b0}^{[t-1]}+\gamma_{b1}^{[t-1]}x_{big}^{[t]})}{1+exp(\gamma_{b0}^{[t-1]}+\gamma_{b1}^{[t-1]}x_{big}^{[t]})} \frac{\Gamma(\phi_{bg}^{[t-1]}+x_{big}^{[t]})(\mu_{big}^{[t-1]})^{x_{big}^{[t]}}}{\Gamma(x_{big}^{[t]})(\phi_{bg}^{[t-1]}+\mu_{big}^{[t-1]})^{\phi_{bg}^{[t-1]}+x_{big}^{[t]}}} & , \text{if } z_{big}^{[t]} = 1. \end{cases}$$

where $\mu_{big}^{[t-1]} = exp(\alpha_g^{[t-1]} + \beta_{gw_{bi}^{[t-1]}}^{[t-1]} + \nu_{bg}^{[t-1]} + \delta_{bi}^{[t-1]})$, and $\Gamma(\cdot)$ represents the Gamma function.

When $z_{big}^{[t]} = 1$, we incorporate a Metropolis-Hasting (MH) step [17]. We sample $x_{big}^*$ from the proposal distribution $NB(\mu_{big}^{[t-1]}, \phi_{bg}^{[t-1]})$ and accept the proposal with probability

$$\rho = \frac{1 + exp(-\gamma_{b0}^{[t-1]} - \gamma_{b1}^{[t-1]}x_{big}^{[t-1]})}{1 + exp(-\gamma_{b0}^{[t-1]} - \gamma_{b1}^{[t-1]}x_{big}^*)}.$$

On the other hand, if $y_{big} > 0$, then $z_{big}^{[t]} = 0$ and $x_{big}^{[t]} = y_{big}$.

2. Update $\gamma_{b0}^{[t]}$ and $\gamma_{b1}^{[t]}$ sequentially. Because

$$L(\gamma_b^{[t]}) \propto \prod_{i=1}^{n_b}\prod_{g=1}^{G} \frac{exp[(\gamma_{b1}^{[t]}x_{big}^{[t]} + \gamma_{b0}^{[t]})z_{big}^{[t]}]}{1 + exp(\gamma_{b1}^{[t]}x_{big}^{[t]} + \gamma_{b0}^{[t]})} \cdot exp(-\frac{(\gamma_{b0}^{[t]})^2}{2\sigma_{z0}^2}) \cdot (-\gamma_{b1}^{[t]})^{a_\gamma - 1} exp(b_\gamma \gamma_{b1}^{[t]}),$$

we update $\gamma_{b0}$ by an MH step with the symmetric proposal distribution $g(\gamma_{b0}^*|\gamma_{b0}^{[t-1]}) \sim N(\gamma_{b0}^{[t-1]}, \sigma_{MH}^2)$. Consequently, the acceptance rate is

$$\rho = \frac{L(\gamma_{b0}^*|-)}{L(\gamma_{b0}^{[t-1]}|-)} = \prod_{i=1}^{n_b}\prod_{g=1}^{G} \frac{exp(\gamma_{b0}^* z_{big}^{[t]})[1 + exp(\gamma_{b1}^{[t-1]}x_{big}^{[t]} + \gamma_{b0}^{[t-1]})]}{exp(\gamma_{b0}^{[t-1]} z_{big}^{[t]})[1 + exp(\gamma_{b1}^{[t-1]}x_{big}^{[t]} + \gamma_{b0}^*)]} \cdot exp(-\frac{(\gamma_{b0}^*)^2 - (\gamma_{b0}^{[t-1]})^2}{2\sigma_{z0}^2}).$$

To update $\gamma_{b1}$, we incorporate an MH step with the proposal distribution $g(-\gamma_{b1}^*|\gamma_{b1}^{[t-1]}) \sim Gamma(-10\gamma_{b1}^{[t-1]}, 10)$, and the acceptance rate being

$$\rho = \frac{L(\gamma_{b1}^*|-)}{L(\gamma_{b1}^{[t-1]}|-)} = \prod_{i=1}^{n_b}\prod_{g=1}^{G} \frac{exp(\gamma_{b1}^* x_{big}^{[t]} z_{big}^{[t]})[1 + exp(\gamma_{b1}^{[t-1]}x_{big}^{[t]} + \gamma_{b0}^{[t]})]}{exp(\gamma_{b1}^{[t-1]} x_{big}^{[t]} z_{big}^{[t]})[1 + exp(\gamma_{b1}^* x_{big}^{[t]} + \gamma_{b0}^{[t]})]}$$
$$\cdot \frac{(-\gamma_{b1}^{[t-1]})^{-a_\gamma - 10\gamma_{b1}^*} 10^{-10\gamma_{b1}^*} \Gamma(-10\gamma_{b1}^{[t-1]})}{(-\gamma_{b1}^*)^{-a_\gamma - 10\gamma_{b1}^{[t-1]}} 10^{-10\gamma_{b1}^{[t-1]}} \Gamma(-10\gamma_{b1}^*)} exp[(10 - b_\gamma)(\gamma_{b1}^{[t-1]} - \gamma_{b1}^*)].$$

3. For each gene $g$, we use an MH step to update $\alpha_g$. Specifically, we let the proposal

distribution be the symmetric $g(\alpha_g^*|\alpha_g^{[t-1]}) \sim N(\alpha_g^{[t-1]}, \sigma_{MH}^2)$ and the acceptance rate be:

$$\rho = \frac{L(\alpha_g^*|-)}{L(\alpha_g^{[t-1]}|-)}$$
$$= \prod_{b=1}^{B}\prod_{i=1}^{n_b} exp((\alpha_g^* - \alpha_g^{[t-1]})x_{big}^{[t]})(\frac{\phi_{bg}^{[t-1]} + exp(\alpha_g^{[t-1]} + \beta_{gw_{bi}^{[t-1]}}^{[t-1]} + \nu_{bg}^{[t-1]} + \delta_{bi}^{[t-1]})}{\phi_{bg}^{[t-1]} + exp(\alpha_g^* + \beta_{gw_{bi}^{[t-1]}}^{[t-1]} + \nu_{bg}^{[t-1]} + \delta_{bi}^{[t-1]})})^{\phi_{bg}^{[t-1]} + x_{big}^{[t]}}$$
$$\cdot exp(-\frac{(\alpha_g^*)^2 - (\alpha_g^{[t-1]})^2}{2\sigma_a^2}).$$

4. For each gene $g$ and for $2 \leq k \leq K$, we sample the indicator $L_{gk}^{[t]}$ from:

$$L_{gk}^{[t]} \sim Bernoulli(\frac{p^{[t-1]}N(\beta_{gk}^{[t-1]}; 0, (\tau_{\beta 1}^{[t-1]})^2)}{p^{[t-1]}N(\beta_{gk}^{[t-1]}; 0, (\tau_{\beta 1}^{[t-1]})^2) + (1 - p^{[t-1]})N(\beta_{gk}^{[t-1]}; 0, \tau_{\beta 0}^2)}).$$

5. Update the inclusion probability $p^{[t]}$ for $L_{gk}^{[t]}$s by sampling:

$$p^{[t]} \sim Beta(\sum_{g=1}^{G}\sum_{k=2}^{K} L_{gk}^{[t]} + a_p, G(K-1) - \sum_{g=1}^{G}\sum_{k=2}^{K} L_{gk}^{[t]} + b_p).$$

6. Update the variance of the spike component of the spike-and-slab prior $(\tau_{\beta 0}^{[t]})^2$ by sampling:

$$(\tau_{\beta 0}^{[t]})^2 \sim Inv - Gamma(a_\tau + \frac{1}{2}\#\{(g,k) : L_{gk}^{[t]} = 0, 1 \leq g \leq G, 2 \leq k \leq K\},$$
$$b_\tau + \frac{1}{2}\sum_{g=1}^{G}\sum_{k=2}^{K} I(L_{gk}^{[t]} = 0) \cdot (\beta_{gk}^{[t-1]})^2),$$

where $\#\{\cdot\}$ represents the number of elements in the set, and $I(\cdot)$ denotes the indicator function.

7. To update $\beta_{gk}^{[t]}$ for cell type two to $K$ and each gene $g$, we use an MH step with the

symmetric proposal distribution $g(\beta^*_{gk}|\beta^{[t-1]}_{gk}) \sim N(\beta^{[t-1]}_{gk}, \sigma^2_{MH})$ and the acceptance rate

$$\rho = \frac{L(\beta^*_{gk}|-)}{L(\beta^{[t-1]}_{gk}|-)}$$
$$= \prod_{(b,i):w^{[t-1]}_{bi}=k} exp((\beta^*_{gk} - \beta^{[t-1]}_{gk})x^{[t]}_{big})(\frac{\phi^{[t-1]}_{bg} + exp(\alpha^{[t]}_g + \beta^{[t-1]}_{gk} + \nu^{[t-1]}_{bg} + \delta^{[t-1]}_{bi})}{\phi^{[t-1]}_{bg} + exp(\alpha^{[t]}_g + \beta^*_{gk} + \nu^{[t-1]}_{bg} + \delta^{[t-1]}_{bi})})^{\phi^{[t-1]}_{bg}+x^{[t]}_{big}}$$
$$\cdot exp(-\frac{(\beta^*_{gk})^2 - (\beta^{[t-1]}_{gk})^2}{2(\tau^{[t]}_{\beta L^{[t]}_{gk}})^2}).$$

8. Update $\nu^{[t]}_{bg}$ by an MH step with the symmetric proposal distribution $g(\nu^*_{bg}|\nu^{[t-1]}_{bg}) \sim N(\nu^{[t-1]}_{bg}, \sigma^2_{MH})$ and the acceptance rate

$$\rho = \frac{L(\nu^*_{bg}|-)}{L(\nu^{[t-1]}_{bg}|-)}$$
$$= \prod_{i=1}^{n_b} exp((\nu^*_{bg} - \nu^{[t-1]}_{bg})x^{[t]}_{big})(\frac{\phi^{[t-1]}_{bg} + exp(\alpha^{[t]}_g + \beta^{[t]}_{gk} + \nu^{[t-1]}_{bg} + \delta^{[t-1]}_{bi})}{\phi^{[t-1]}_{bg} + exp(\alpha^{[t]}_g + \beta^{[t]}_{gk} + \nu^*_{bg} + \delta^{[t-1]}_{bi})})^{\phi^{[t-1]}_{bg}+x^{[t]}_{big}}$$
$$\cdot exp(-\frac{(\nu^*_{bg})^2 - (\nu^{[t-1]}_{bg})^2}{2\sigma^2_c}).$$

9. Update $\delta^{[t]}_{bi}$ by an MH step with the symmetric proposal distribution $g(\delta^*_{bi}|\delta^{[t-1]}_{bi}) \sim N(\delta^{[t-1]}_{bi}, \sigma^2_{MH})$ and the acceptance rate

$$\rho = \frac{L(\delta^*_{bi}|-)}{L(\delta^{[t-1]}_{bi}|-)}$$
$$= \prod_{g=1}^{G} exp((\delta^*_{bi} - \delta^{[t-1]}_{bi})x^{[t]}_{big})(\frac{\phi^{[t-1]}_{bg} + exp(\alpha^{[t]}_g + \beta^{[t]}_{gk} + \nu^{[t]}_{bg} + \delta^{[t-1]}_{bi})}{\phi^{[t-1]}_{bg} + exp(\alpha^{[t]}_g + \beta^{[t]}_{gk} + \nu^{[t]}_{bg} + \delta^*_{bi})})^{\phi^{[t-1]}_{bg}+x^{[t]}_{big}}$$
$$\cdot exp(-\frac{(\delta^*_{bi})^2 - (\delta^{[t-1]}_{bi})^2}{2\sigma^2_d}).$$

10. Update $\phi^{[t]}_{bg}$ by an MH step with the proposal distribution $g(\phi^*_{bg}|\phi^{[t-1]}_{bg}) \sim Gamma(\phi^{[t-1]}_{bg}, 1)$

and the acceptance rate

$$\rho = \frac{L(\phi_{bg}^{*})g(\phi_{bg}^{[t-1]}|\phi_{bg}^{*})}{L(\phi_{bg}^{[t-1]})g(\phi_{bg}^{*}|\phi_{bg}^{[t-1]})}$$

$$= \prod_{i=1}^{n_b}[\frac{\Gamma(\phi_{bg}^{*}+x_{big}^{[t]})(\phi_{bg}^{*})^{\phi_{bg}^{*}}}{\Gamma(\phi_{bg}^{*})(\phi_{bg}^{*}+\eta_{big}^{[t]})^{\phi_{bg}^{*}+x_{big}^{[t]}}} \cdot \frac{\Gamma(\phi_{bg}^{[t-1]})(\phi_{bg}^{[t-1]}+\eta_{big}^{[t]})^{\phi_{bg}^{[t-1]}+x_{big}^{[t]}}}{\Gamma(\phi_{bg}^{[t-1]}+x_{big}^{[t]})(\phi_{bg}^{[t-1]})^{\phi_{bg}^{[t-1]}}}]$$

$$\cdot \frac{(\phi_{bg}^{*})^{\kappa-1}}{(\phi_{bg}^{[t-1]})^{\kappa-1}} exp(-\tau(\phi_{bg}^{*}-\phi_{bg}^{[t-1]})) \frac{(\phi_{bg}^{[t-1]})^{\phi_{bg}^{*}-1}\Gamma(\phi_{bg}^{[t-1]})}{(\phi_{bg}^{*})^{\phi_{bg}^{[t-1]}-1}\Gamma(\phi_{bg}^{*})} exp(\phi_{bg}^{*}-\phi_{bg}^{[t-1]}),$$

where $\eta_{big}^{[t]} = exp(\alpha_g^{[t]} + \beta_{gw_{bi}^{[t-1]}}^{[t]} + \nu_{bg}^{[t]} + \delta_{bi}^{[t]})$ denotes the mean gene expression level for gene $g$ in cell $i$ of batch $b$.

11. The conditional posterior distribution for the cell type indicator $w_{bi}^{[t]}$ of cell $i$ in batch $b$ is:

$$Pr(w_{bi}^{[t]} = k|-) \propto \pi_{bk}^{[t-1]} \prod_{g=1}^{G} \frac{exp[(\alpha_g^{[t]} + \beta_{gk}^{[t]} + \nu_{bg}^{[t]} + \delta_{bi}^{[t]})x_{big}^{[t]}]}{(exp(\alpha_g^{[t]} + \beta_{gk}^{[t]} + \nu_{bg}^{[t]} + \delta_{bi}^{[t]}) + \phi_{bg}^{[t]})^{x_{big}^{[t]}+\phi_{bg}^{[t]}}}.$$

We implement an MH step with the symmetric proposal distribution $Pr(w_{bi}^{[t]} = k^*|w_{bi}^{[t-1]} = k) \sim Multinomial(1; \frac{1}{K}, \cdots, \frac{1}{K})$ and the acceptance rate

$$\rho = \frac{Pr(w_{bi}^{[t]} = k^*|-)}{Pr(w_{bi}^{[t]} = k|-)}$$

$$= \frac{\pi_{bk^*}^{[t-1]}}{\pi_{bk}^{[t-1]}} \prod_{g=1}^{G} exp[(\beta_{gk^*}^{[t]} - \beta_{gk}^{[t]})x_{big}^{[t]}](\frac{exp(\alpha_g^{[t]} + \beta_{gk}^{[t]} + \nu_{bg}^{[t]} + \delta_{bi}^{[t]}) + \phi_{bg}^{[t]}}{exp(\alpha_g^{[t]} + \beta_{gk^*}^{[t]} + \nu_{bg}^{[t]} + \delta_{bi}^{[t]}) + \phi_{bg}^{[t]}})^{x_{big}^{[t]}+\phi_{bg}^{[t]}}.$$

12. Update $\pi_b^{[t]}$ by sampling from the Dirichlet distribution

$$Dir(\xi + \sum_{i=1}^{n_b} 1(w_{bi}^{[t]} = 1), \xi + \sum_{i=1}^{n_b} 1(w_{bi}^{[t]} = 2), \cdots, \xi + \sum_{i=1}^{n_b} 1(w_{bi}^{[t]} = K)).$$

The Markov chain of the MCMC algorithm can get stuck in the local modes of the posterior distribution for a long period of time. In principle, we can further incorporate the Metropolis coupled MCMC algorithm [18] to jump out of the local modes more easily. In practice, we recommend running multiple chains with different initial values and then choosing the chain that gives the largest value of the observed data likelihood to conduct the posterior inference. According to our experiences, we can usually achieve good posterior estimations with five Markov chains each with a different initial value by randomly sampling a seed from 1 to 10,000.

#### Selecting the total number of cell types

If the total number of cell types $K$ is unknown, we apply Bayesian information criterion (BIC) [19] to select the optimal $K^*$ for the minimum BIC value based on the observed likelihood $L_o(\widehat{\Theta}|\boldsymbol{y})$ as Eq. 2.

$$BIC(K) = -2L_o(\widehat{\Theta}|\boldsymbol{y}) + [K(B+G) + 2B + (2B-1)G + \sum_{b=1}^{B}(n_b - 1)] \cdot log(\sum_{b=1}^{B} n_b G).$$

### Theorems and Proofs

**Lemma 1.** *Let $\mathcal{F}^G$ be the family of $G(\geq 2)$-dimensional multivariate distribution with the probability mass function for $\boldsymbol{y} = (y_1, \cdots, y_G)$ as*

$$f^G(\boldsymbol{y}|\gamma, \boldsymbol{\phi}, \boldsymbol{\mu}) = \prod_{g=1}^{G}\{[\frac{1}{1+exp(\gamma_0+\gamma_1 y_g)} f_{NB}(y_g;\mu_g,\phi_g)]^{1(y_g>0)} \cdot [\sum_{x=1}^{\infty}\frac{exp(\gamma_0+\gamma_1 x)}{1+exp(\gamma_0+\gamma_1 x)} f_{NB}(x;\mu_g,\phi_g) + f_{NB}(0;\mu_g,\phi_g)]^{1(y_g=0)}\} \tag{4}$$

*such that $\gamma_1 < 0$ and for any two distinct elements $f^G_k(\boldsymbol{y}) = f^G(\boldsymbol{y}|\gamma, \boldsymbol{\phi}, \boldsymbol{\mu}_k) \in \mathcal{F}^G, k = 1, 2,$ there exist at least two dimensions $l$ and $m$ with $\mu_{l1} \neq \mu_{l2}$ and $\mu_{m1} \neq \mu_{m2}$, then the class of all finite mixtures*

$$\mathcal{H}^G = \{h^G(\boldsymbol{y}|\gamma, \boldsymbol{\phi}, \boldsymbol{\mu}_1, \boldsymbol{\mu}_2, \cdots, \boldsymbol{\mu}_K) | h^G(\boldsymbol{y}|\gamma, \boldsymbol{\phi}, \boldsymbol{\mu}_1, \boldsymbol{\mu}_2, \cdots, \boldsymbol{\mu}_K) = \sum_{k=1}^{K}\pi_k f^G(\boldsymbol{y}|\gamma, \boldsymbol{\phi}, \boldsymbol{\mu}_k), f^G(\boldsymbol{y}|\gamma, \boldsymbol{\phi}, \boldsymbol{\mu}_k) \in \mathcal{F}^G, \pi_k > 0, k = 1, 2, \cdots, K, \sum_{k=1}^{K}\pi_k = 1\}$$

*is identifiable (up to label switching).*

*Proof.* We reparameterize $(\mu_g, \phi_g)$ as $(p_g, \phi_g)$ such that $p_g = \frac{\mu_g}{\mu_g+\phi_g}$ for all $g = 1, 2, \cdots, G$. Consequently, the identifiability with respect to $(\boldsymbol{\gamma}, \boldsymbol{\phi}, \boldsymbol{\mu})$ is equivalent to that with respect to $(\boldsymbol{\gamma}, \boldsymbol{\phi}, \boldsymbol{p})$. With a little bit abuse of notations, we still use $f^G(\boldsymbol{y}|\boldsymbol{\gamma}, \boldsymbol{\phi}, \boldsymbol{p})$ to indicate the probability mass function of the new parameterization hereafter. Suppose that the finite mixture of $\mathcal{F}^G$ is not identifiable, then we have two different representations of the probability mass function $h(\boldsymbol{y})$ of the same finite mixtures:

$$h(\boldsymbol{y}) = \sum_{k=1}^{K}\pi_k f^G(\boldsymbol{y}|\boldsymbol{\gamma}, \boldsymbol{\phi}, \boldsymbol{p}_k) = \sum_{l=1}^{L}\xi_l f^G(\boldsymbol{y}|\boldsymbol{\delta}, \boldsymbol{\psi}, \boldsymbol{r}_l). \tag{5}$$

where the tuples $(\boldsymbol{\gamma}, \boldsymbol{\phi}, \boldsymbol{p}_k)$ for $k = 1, 2, \cdots, K$ are mutually distinct, and so are the tuples $(\boldsymbol{\delta}, \boldsymbol{\psi}, \boldsymbol{r}_l)$ for $l = 1, 2, \cdots, L$.

We define a total ordering ($\succeq$) of $\mathcal{F}^G$. For $f_1^G, f_2^G \in \mathcal{F}^G$, $f_1^G \succeq f_2^G$ if:

1. there exists a $g \geq 1$ such that for all $j < g$, $p_{j1} = p_{j2}$ and $\phi_{j1} = \phi_{j2}$ but $p_{g1} > p_{g2}$;
2. or there exists a $g$ such that for all $j < g$, $p_{j1} = p_{j2}$ and $\phi_{j1} = \phi_{j2}$ as well as $p_{g1} = p_{g2}$ but $\phi_{g1} > \phi_{g2}$;
3. or $\boldsymbol{p}_1 = \boldsymbol{p}_2$ and $\boldsymbol{\phi}_1 = \boldsymbol{\phi}_2$ but $\gamma_{11} < \gamma_{21}$ ;
4. or $\boldsymbol{p}_1 = \boldsymbol{p}_2, \boldsymbol{\phi}_1 = \boldsymbol{\phi}_2$ and $\gamma_{11} = \gamma_{21}$ but $\gamma_{10} \leq \gamma_{20}$.

Without loss of generality, we assume that $f^G(\boldsymbol{y}|\boldsymbol{\gamma}, \boldsymbol{\phi}, \boldsymbol{p}_1) \succeq f^G(\boldsymbol{y}|\boldsymbol{\delta}, \boldsymbol{\psi}, \boldsymbol{r}_1)$ and the mixture components on both sides of (5) are ordered:

$$f^G(\boldsymbol{y}|\boldsymbol{\gamma}, \boldsymbol{\phi}, \boldsymbol{p}_1) \succeq f^G(\boldsymbol{y}|\boldsymbol{\gamma}, \boldsymbol{\phi}, \boldsymbol{p}_2) \succeq \cdots \succeq f^G(\boldsymbol{y}|\boldsymbol{\gamma}, \boldsymbol{\phi}, \boldsymbol{p}_K),$$
$$f^G(\boldsymbol{y}|\boldsymbol{\delta}, \boldsymbol{\psi}, \boldsymbol{r}_1) \succeq f^G(\boldsymbol{y}|\boldsymbol{\delta}, \boldsymbol{\psi}, \boldsymbol{r}_2) \succeq \cdots \succeq f^G(\boldsymbol{y}|\boldsymbol{\delta}, \boldsymbol{\psi}, \boldsymbol{r}_L).$$

**For $k = 1$, we use mathematical induction to prove that for every $G_0 \in \{1, 2, \cdots, G\}$,**

$$r_{j1} = p_{j1}, \phi_j = \psi_j, \forall j \in \{1, 2, \cdots, G_0\}, \tag{*}$$

**and there exist a $K_{G_0}$ and an $L_{G_0}$ such that**

$$\sum_{k=1}^{K_{G_0}} \pi_k f^{G-G_0}(\boldsymbol{y}_{-G_0}|\boldsymbol{\gamma}, \boldsymbol{\phi}_{-G_0}, \boldsymbol{p}_{-G_0,k}) = \sum_{l=1}^{L_{G_0}} \xi_l f^{G-G_0}(\boldsymbol{y}_{-G_0}|\boldsymbol{\delta}, \boldsymbol{\psi}_{-G_0}, \boldsymbol{r}_{-G_0,l}), \tag{**}$$

**where the subscript $-G_0$ denotes that the first $G_0$ entries in the original vectors are excluded. Specifically, $\boldsymbol{y}_{-G_0} = (y_{G_0+1}, y_{G_0+2}, \cdots, y_G)^T$.**

We first prove Equations (*) and (**) hold for $G_0 = 1$. We define a linear mapping that maps a probability distribution of $\mathcal{F}^G$ to a function that shares a similar spirit as a probability generating function $M_1 : f^G(\boldsymbol{y}) \in \mathcal{F}^G \to \Phi_1(t_1, \boldsymbol{y}_{-1}) \in \mathcal{G}_1$ such that $M_1(f^G(\boldsymbol{y})) = \Phi_1(t_1, \boldsymbol{y}_{-1}) = \sum_{y_1=1}^{\infty} f^G(\boldsymbol{y}|\boldsymbol{\gamma}, \boldsymbol{\phi}, \boldsymbol{p}) t_1^{y_1} = \sum_{y_1=1}^{\infty} f^1(y_1|\boldsymbol{\gamma}, \phi_1, p_1) t_1^{y_1} \cdot f^{G-1}(\boldsymbol{y}_{-1}|\boldsymbol{\gamma}, \boldsymbol{\phi}_{-1}, \boldsymbol{p}_{-1})$. Notice that $\Phi_1(t_1, \boldsymbol{y}_{-1})$ does not include the term of $y_1 = 0$, that is, $f^1(0|\boldsymbol{\gamma}, \phi_1, p_1) t_1^0 \cdot f^{G-1}(\boldsymbol{y}_{-1}|\boldsymbol{\gamma}, \boldsymbol{\phi}_{-1}, \boldsymbol{p}_{-1})$. Specifically, we denote $\Phi_{1k}(t_1, \boldsymbol{y}_{-1}) = M_1(f^G(\boldsymbol{y}|\boldsymbol{\gamma}, \boldsymbol{\phi}, \boldsymbol{p}_k))$ and $\Psi_{1l}(t_1, \boldsymbol{y}_{-1}) = M_1(f^G(\boldsymbol{y}|\boldsymbol{\delta}, \boldsymbol{\psi}, \boldsymbol{r}_l)) \in \mathcal{G}_1$ for $k = 1, 2, \cdots, K$ and $l = 1, 2, \cdots, L$. It is noteworthy that $M_1$ is a linear mapping so that if applying $M_1$ to both sides of Equation (5), then we have

$$\sum_{k=1}^{K} \pi_k \Phi_{1k}(t_1, \boldsymbol{y}_{-1}) = \sum_{l=1}^{L} \xi_l \Psi_{1l}(t_1, \boldsymbol{y}_{-1}). \tag{6}$$

More specifically,

$$\Phi_{1k}(t_1, \boldsymbol{y}_{-1}) = \sum_{y_1=1}^{\infty} \frac{1}{1+exp(\gamma_0+\gamma_1 y_1)} C_{y_1}^{\phi_1+y_1-1}(p_{1k})^{y_1}(1-p_{1k})^{\phi_1}(t_1)^{y_1} \cdot f^{G-1}(\boldsymbol{y}_{-1}|\boldsymbol{\gamma}, \boldsymbol{\phi}_{-1}, \boldsymbol{p}_{-1,k})$$
$$= [(\frac{1-p_{1k}}{1-p_{1k}t_1})^{\phi_1} - R_{1k}(t_1)] \cdot f^{G-1}(\boldsymbol{y}_{-1}|\boldsymbol{\gamma}, \boldsymbol{\phi}_{-1}, \boldsymbol{p}_{-1,k}), \quad (7)$$

where $R_{1k}(t_1) = (1-p_{1k})^{\phi_1} + \sum_{y_1=1}^{\infty} \frac{exp(\gamma_0+\gamma_1 y_1)}{1+exp(\gamma_0+\gamma_1 y_1)} C_{y_1}^{\phi_1+y_1-1}(p_{1k}t_1)^{y_1}(1-p_{1k})^{\phi_1}$ is the residual part. Let $t_1 \to \frac{1}{p_{11}}$, because $\gamma_1 < 0$ so that $\frac{p_{1k}exp(\gamma_1)}{p_{11}} < \frac{p_{1k}}{p_{11}} \le 1$, we have

$$\lim_{t_1 \to \frac{1}{p_{11}}} R_{1k}(t_1) \le (1-p_{1k})^{\phi_1} + \lim_{t_1 \to \frac{1}{p_{11}}} \sum_{y_1=1}^{\infty} exp(\gamma_0+\gamma_1 y_1) C_{y_1}^{\phi_1+y_1-1}(p_{1k}t_1)^{y_1}(1-p_{1k})^{\phi_1}$$
$$= [1-exp(\gamma_0)](1-p_{1k})^{\phi_1} + exp(\gamma_0)(\frac{1-p_{1k}}{1-p_{1k}exp(\gamma_1)/p_{11}})^{\phi_1} < \infty, \quad (8)$$

Similarly, $\Psi_{1l}(t_1, \boldsymbol{y}_{-1}) = \sum_{y_1=1}^{\infty} f^G(\boldsymbol{y}|\boldsymbol{\delta}, \boldsymbol{\psi}, \boldsymbol{r}_l)t_1^{y_1} = [(\frac{1-r_{1l}}{1-r_{1l}t_1})^{\psi_1} - S_{1l}(t_1)] \cdot f^{G-1}(\boldsymbol{y}_{-1}|\boldsymbol{\delta}, \boldsymbol{\psi}_{-1}, \boldsymbol{r}_{-1,l})$,

$$S_{1l}(t_1) = (1-r_{1l})^{\psi_1} + \sum_{y_1=1}^{\infty} exp(\delta_0+\delta_1 y_1) C_{y_1}^{\psi_1+y_1-1}(r_{1l}t_1)^{y_1}(1-r_{1l})^{\psi_1}$$
$$\le [1-exp(\delta_0)](1-r_{1l})^{\psi_1} + exp(\delta_0)(\frac{1-r_{1l}}{1-r_{1l}exp(\delta_1)t_1})^{\psi_1} \quad (9)$$

As $t_1 \to \frac{1}{p_{11}}$, because $\frac{r_{1l}exp(\gamma_1)}{p_{11}} < \frac{r_{1l}}{p_{11}} \le \frac{r_{11}}{p_{11}} \le 1$, we have $\lim_{t_1 \to \frac{1}{p_{11}}} S_{1l}(t_1) < \infty$.

Notice that $f^G(\boldsymbol{y}|\boldsymbol{\gamma}, \boldsymbol{\phi}, \boldsymbol{p}_1) \succeq f^G(\boldsymbol{y}|\boldsymbol{\delta}, \boldsymbol{\psi}, \boldsymbol{r}_1)$ implies $r_{11} < p_{11}$ or $r_{11} = p_{11}, \psi_1 \le \phi_1$. According to (8) and (9), we have

$$\lim_{t_1 \to \frac{1}{p_{11}}} \frac{\Psi_{1l}(t_1, \boldsymbol{y}_{-1})}{\Phi_{11}(t_1, \boldsymbol{y}_{-1})} = \lim_{t_1 \to \frac{1}{p_{11}}} \frac{(\frac{1-r_{1l}}{1-r_{1l}t_1})^{\psi_1} - S_{1l}(t_1)}{(\frac{1-p_{11}}{1-p_{11}t_1})^{\phi_1} - R_{11}(t_1)} \cdot \frac{f^{G-1}(\boldsymbol{y}_{-1}|\boldsymbol{\delta}, \boldsymbol{\psi}_{-1}, \boldsymbol{r}_{-1,l})}{f^{G-1}(\boldsymbol{y}_{-1}|\boldsymbol{\gamma}, \boldsymbol{\phi}_{-1}, \boldsymbol{p}_{-1,1})}$$
$$= \frac{f^{G-1}(\boldsymbol{y}_{-1}|\boldsymbol{\delta}, \boldsymbol{\psi}_{-1}, \boldsymbol{r}_{-1,l})}{f^{G-1}(\boldsymbol{y}_{-1}|\boldsymbol{\gamma}, \boldsymbol{\phi}_{-1}, \boldsymbol{p}_{-1,1})} \cdot \lim_{t_1 \to \frac{1}{p_{11}}} \frac{(\frac{1-r_{1l}}{1-r_{1l}t_1})^{\psi_1}(1-p_{11}t_1)^{\phi_1} - S_{1l}(t_1)(1-p_{11}t_1)^{\phi_1}}{(1-p_{11})^{\phi_1} - R_{11}(t_1)(1-p_{11}t_1)^{\phi_1}}$$
$$= \begin{cases} \frac{f^{G-1}(\boldsymbol{y}_{-1}|\boldsymbol{\delta}, \boldsymbol{\psi}_{-1}, \boldsymbol{r}_{-1,l})}{f^{G-1}(\boldsymbol{y}_{-1}|\boldsymbol{\gamma}, \boldsymbol{\phi}_{-1}, \boldsymbol{p}_{-1,1})} & , \ if \ r_{1l} = p_{11}, \psi_1 = \phi_1 \\ 0 & , \ if \ r_{1l} = p_{11}, \psi_1 < \phi_1 \\ 0 & , \ if \ r_{1l} < p_{11} \end{cases} \quad (10)$$

If $r_{11} < p_{11}$ or $r_{11} = p_{11}, \psi_1 < \phi_1$, then dividing $\Phi_{11}(t_1, \boldsymbol{y}_{-1})$ on both sides of Equation (6) and let $t_1 \to \frac{1}{p_{11}}$, we have

$$\lim_{t_1 \to \frac{1}{p_{11}}} \sum_{k=1}^{K} \pi_k \frac{\Phi_{1k}(t_1, \boldsymbol{y}_{-1})}{\Phi_{11}(t_1, \boldsymbol{y}_{-1})} \geq \lim_{t_1 \to \frac{1}{p_{11}}} \pi_1 \frac{\Phi_{11}(t_1, \boldsymbol{y}_{-1})}{\Phi_{11}(t_1, \boldsymbol{y}_{-1})} = \pi_1 > 0 = \lim_{t_1 \to \frac{1}{p_{11}}} \sum_{l=1}^{L} \xi_l \frac{\Psi_{1l}(t_1, \boldsymbol{y}_{-1})}{\Phi_{11}(t_1, \boldsymbol{y}_{-1})},$$

which contradicts with Equation (5). Thus, $r_{11} = p_{11}$ and $\psi_1 = \phi_1$, which means Equation (*) holds. Similar to Equation (10), we have

$$\lim_{t_1 \to \frac{1}{p_{11}}} \frac{\Phi_{1k}(t_1, \boldsymbol{y}_{-1})}{\Phi_{11}(t_1, \boldsymbol{y}_{-1})} = \begin{cases} \frac{f^{G-1}(\boldsymbol{y}_{-1}|\boldsymbol{\gamma}, \boldsymbol{\phi}_{-1}, \boldsymbol{p}_{-1,k})}{f^{G-1}(\boldsymbol{y}_{-1}|\boldsymbol{\gamma}, \boldsymbol{\phi}_{-1}, \boldsymbol{p}_{-1,1})} & , \ if \ p_{1k} = p_{11} \\ 0 & , \ if \ p_{1k} < p_{11} \end{cases}$$

Moreover, there exists a $K_1 \leq K$ such that $p_{1k} = p_{11}$ for $k = 1, 2, \cdots, K_1$ but $p_{1k} < p_{11}$ for $k = K_1 + 1, \cdots, K$. There also exists an $L_1 \leq L$ such that $r_{1l} = p_{11}$ for $l = 1, 2, \cdots, L_1$ but $r_{1l} < p_{11}$ for $l = L_1 + 1, \cdots, L$. $p_{11} = p_{11}$ and $r_{11} = p_{11}$, therefore, $K_1 \geq 1$ and $L_1 \geq 1$. Dividing $\Phi_{11}(t_1, \boldsymbol{y}_{-1})$ on both sides of Equation (6) and let $t_1 \to \frac{1}{p_{11}}$, we have

$$\begin{aligned} \sum_{k=1}^{K_1} \pi_k \frac{\Phi_{1k}(t_1, \boldsymbol{y}_{-1})}{\Phi_{11}(t_1, \boldsymbol{y}_{-1})} &= \sum_{l=1}^{L_1} \xi_l \frac{\Psi_{1l}(t_1, \boldsymbol{y}_{-1})}{\Phi_{11}(t_1, \boldsymbol{y}_{-1})} \\ \Rightarrow \sum_{k=1}^{K_1} \pi_k f^{G-1}(\boldsymbol{y}_{-1}|\boldsymbol{\gamma}, \boldsymbol{\phi}_{-1}, \boldsymbol{p}_{-1,k}) &= \sum_{l=1}^{L_1} \xi_l f^{G-1}(\boldsymbol{y}_{-1}|\boldsymbol{\delta}, \boldsymbol{\psi}_{-1}, \boldsymbol{r}_{-1,l}). \end{aligned} \tag{11}$$

Thus, we have proven that Equation (**) holds.

Now let us assume that Equations (*) and (**) hold for for $G_0 = g$. In other words, $p_{j1} = r_{j1}, \phi_j = \psi_j$ for all $j = 1, 2, \cdots, g$ and there are $K_g \geq 1$ and $L_g \geq 1$ such that

$$\sum_{k=1}^{K_g} \pi_k f^{G-g}(\boldsymbol{y}_{-g}|\boldsymbol{\gamma}, \boldsymbol{\phi}_{-g}, \boldsymbol{p}_{-g,k}) = \sum_{l=1}^{L_g} \xi_l f^{G-g}(\boldsymbol{y}_{-g}|\boldsymbol{\delta}, \boldsymbol{\psi}_{-g}, \boldsymbol{r}_{-g,l}). \tag{12}$$

Let $G_0 = g + 1$. Similar to $M_1$, we define a linear map $M_{g+1} : \mathcal{F}^{G-g} \to \mathcal{G}_{g+1}$ such that $M_{g+1}(f^{G-g}(\boldsymbol{y}_{-g}|\boldsymbol{\gamma}, \boldsymbol{\phi}_{-g}, \boldsymbol{p}_{-g})) = \Phi_{g+1}(t_{g+1}, \boldsymbol{y}_{-(g+1)}) = \sum_{y_{g+1}=1}^{\infty} f^{G-g}(\boldsymbol{y}_{-g}|\boldsymbol{\gamma}, \boldsymbol{\phi}_{-g}, \boldsymbol{p}_{-g}) t_{g+1}^{y_{g+1}} =$

$\sum_{y_{g+1}=1}^{\infty} f^1(y_{g+1}|\gamma, \phi_{g+1}, p_{g+1}) t_{g+1}^{y_{g+1}} \cdot f^{G-(g+1)}(\boldsymbol{y}_{-(g+1)}|\boldsymbol{\gamma}, \boldsymbol{\phi}_{-(g+1)}, \boldsymbol{p}_{-(g+1)})$. Consequently,

$$\begin{aligned} &M_{g+1}(f^{G-g}(\boldsymbol{y}_{-g}|\boldsymbol{\gamma}, \boldsymbol{\phi}_{-g}, \boldsymbol{p}_{-g,k})) = \Phi_{g+1,k}(t_{g+1}, \boldsymbol{y}_{-(g+1)}) \\ &= [(\frac{1 - p_{g+1,k}}{1 - p_{g+1,k} t_{g+1}})^{\phi_{g+1}} - R_{g+1,k}(t_{g+1})] \cdot f^{G-(g+1)}(\boldsymbol{y}_{-(g+1)}|\boldsymbol{\gamma}, \boldsymbol{\phi}_{-(g+1)}, \boldsymbol{p}_{-(g+1),k}), \ k = 1, \cdots, K_g; \end{aligned}$$

$$M_{g+1}(f^{G-g}(\boldsymbol{y}_{-g}|\boldsymbol{\delta},\boldsymbol{\psi}_{-g},\boldsymbol{r}_{-g,l})) = \Psi_{g+1,l}(t_{g+1},\boldsymbol{y}_{-(g+1)})$$
$$= [(\frac{1-r_{g+1,l}}{1-p_{g+1,l}t_{g+1}})^{\psi_{g+1}} - S_{g+1,l}(t_{g+1})]\cdot f^{G-(g+1)}(\boldsymbol{y}_{-(g+1)}|\boldsymbol{\delta},\boldsymbol{\psi}_{-(g+1)},\boldsymbol{r}_{-(g+1),l}),\ l=1,\cdots,L_g.$$

If we apply $M_{g+1}$ to both sides of Equation (12), then we have

$$\sum_{k=1}^{K_g}\pi_k\Phi_{g+1,k}(t_{g+1},\boldsymbol{y}_{-(g+1)}) = \sum_{l=1}^{L_g}\xi_l\Psi_{g+1,l}(t_{g+1},\boldsymbol{y}_{-(g+1)}). \tag{13}$$

Notice that given $p_{j1}=r_{j1},\phi_j=\psi_j$ for all $j=1,2,\cdots,g$, $f^G(\boldsymbol{y}|\boldsymbol{\gamma},\boldsymbol{\phi},\boldsymbol{p}_1)\succeq f^G(\boldsymbol{y}|\boldsymbol{\delta},\boldsymbol{\psi},\boldsymbol{r}_1)$ implies $r_{g+1,1}<p_{g+1,1}$ or $r_{g+1,1}=p_{g+1,1},\psi_{g+1}\le\phi_{g+1}$. Similar to Equation (10), we have

$$\begin{aligned}
&\lim_{t_{g+1}\to\frac{1}{p_{g+1,1}}}\frac{\Psi_{g+1,l}(t_{g+1},\boldsymbol{y}_{-(g+1)})}{\Phi_{g+1,1}(t_{g+1},\boldsymbol{y}_{-(g+1)})}\\
&=\lim_{t_{g+1}\to\frac{1}{p_{g+1,1}}}\frac{(\frac{1-r_{g+1,l}}{1-r_{g+1,l}t_{g+1}})^{\psi_{g+1}}-S_{g+1,l}(t_{g+1})}{(\frac{1-p_{g+1,1}}{1-p_{g+1,1}t_{g+1}})^{\phi_{g+1}}-R_{g+1,1}(t_{g+1})}\cdot\frac{f^{G-(g+1)}(\boldsymbol{y}_{-(g+1)}|\boldsymbol{\delta},\boldsymbol{\psi}_{-(g+1)},\boldsymbol{r}_{-(g+1),l})}{f^{G-(g+1)}(\boldsymbol{y}_{-(g+1)}|\boldsymbol{\gamma},\boldsymbol{\phi}_{-(g+1)},\boldsymbol{p}_{-(g+1),1})}\\
&=\frac{f^{G-(g+1)}(\boldsymbol{y}_{-(g+1)}|\boldsymbol{\delta},\boldsymbol{\psi}_{-(g+1)},\boldsymbol{r}_{-(g+1),l})}{f^{G-(g+1)}(\boldsymbol{y}_{-(g+1)}|\boldsymbol{\gamma},\boldsymbol{\phi}_{-(g+1)},\boldsymbol{p}_{-(g+1),1})}\cdot\lim_{t_{g+1}\to\frac{1}{p_{g+1,1}}}\frac{(\frac{1-r_{g+1,l}}{1-r_{g+1,l}t_{g+1}})^{\psi_{g+1}}-S_{g+1,l}(t_{g+1})}{(\frac{1-p_{g+1,1}}{1-p_{g+1,1}t_{g+1}})^{\phi_{g+1}}-R_{g+1,1}(t_{g+1})}\\
&=\begin{cases}\frac{f^{G-(g+1)}(\boldsymbol{y}_{-(g+1)}|\boldsymbol{\delta},\boldsymbol{\psi}_{-(g+1)},\boldsymbol{r}_{-(g+1),l})}{f^{G-(g+1)}(\boldsymbol{y}_{-(g+1)}|\boldsymbol{\gamma},\boldsymbol{\phi}_{-(g+1)},\boldsymbol{p}_{-(g+1),1})} & ,\ if\ r_{g+1,l}=p_{g+1,1},\psi_{g+1}=\phi_{g+1}\\ 0 & ,\ if\ r_{g+1,l}=p_{g+1,1},\psi_{g+1}<\phi_{g+1}\\ 0 & ,\ if\ r_{g+1,l}<p_{g+1,1}\end{cases}
\end{aligned}\tag{14}$$

If $r_{g+1,l}<p_{g+1,1}$ or $r_{g+1,l}=p_{g+1,1},\psi_{g+1}<\phi_{g+1}$, then dividing $\Phi_{g+1,1}(t_{g+1},\boldsymbol{y}_{-(g+1)})$ on both sides of Equation (13) and letting $t_{g+1}\to\frac{1}{p_{g+1,1}}$, we have

$$\begin{aligned}
\lim_{t_{g+1}\to\frac{1}{p_{g+1,1}}}\sum_{k=1}^{K_g}\pi_k\frac{\Phi_{g+1,k}(t_{g+1},\boldsymbol{y}_{-(g+1)})}{\Phi_{g+1,1}(t_{g+1},\boldsymbol{y}_{-(g+1)})}&\ge\lim_{t_{g+1}\to\frac{1}{p_{g+1,1}}}\pi_1\frac{\Phi_{g+1,1}(t_{g+1},\boldsymbol{y}_{-(g+1)})}{\Phi_{g+1,1}(t_{g+1},\boldsymbol{y}_{-(g+1)})}=\pi_1\\
&>0=\lim_{t_{g+1}\to\frac{1}{p_{g+1,1}}}\sum_{l=1}^{L_g}\xi_l\frac{\Psi_{g+1,l}(t_{g+1},\boldsymbol{y}_{-(g+1)})}{\Phi_{g+1,1}(t_{g+1},\boldsymbol{y}_{-(g+1)})},
\end{aligned}\tag{15}$$

which contradicts with (12). Thus, $r_{g+1,l}=p_{g+1,1}$ and $\psi_{g+1}=\phi_{g+1}$, which means that Equation $(*)$ holds. Similar to Equation (14), for $k=1,2,\cdots,K_g$, we have

$$\lim_{t_{g+1}\to\frac{1}{p_{g+1,1}}}\frac{\Phi_{g+1,k}(t_{g+1},\boldsymbol{y}_{-(g+1)})}{\Phi_{g+1,1}(t_{g+1},\boldsymbol{y}_{-(g+1)})}=\begin{cases}\frac{f^{G-(g+1)}(\boldsymbol{y}_{-(g+1)}|\boldsymbol{\gamma},\boldsymbol{\phi}_{-(g+1)},\boldsymbol{p}_{-(g+1),k})}{f^{G-(g+1)}(\boldsymbol{y}_{-(g+1)}|\boldsymbol{\gamma},\boldsymbol{\phi}_{-(g+1)},\boldsymbol{p}_{-(g+1),1})} & ,\ if\ p_{g+1,k}=p_{g+1,1}\\ 0 & ,\ if\ p_{g+1,k}<p_{g+1,1}\end{cases}$$

Further, there exists a $K_{g+1}\le K_g$ such that $p_{g+1,k}=p_{g+1,1}$ for $k=1,2,\cdots,K_{g+1}$ but

$p_{g+1,k} < p_{g+1,1}$ for $k = K_{g+1}+1, K_{g+1}+2, \cdots, K_g$. There also exists an $L_{g+1} \leq L_g$ such that $r_{g+1,l} = p_{g+1,1}$ for $l = 1, 2, \cdots, L_{g+1}$ but $r_{g+1,l} < p_{g+1,1}$ for $l = L_{g+1}+1, L_{g+1}+2, \cdots, L_g$. $p_{g+1,1} = p_{g+1,1}$ and $p_{g+1,1} = r_{g+1,1}$, therefore, $K_{g+1} \geq 1$ and $L_{g+1} \geq 1$. Dividing $\Phi_{g+1,1}(t_{g+1}, \boldsymbol{y}_{-(g+1)})$ on both sides of Equation (13) and letting $t_{g+1} \to \frac{1}{p_{g+1,1}}$, we have,

$$\sum_{k=1}^{K_{g+1}} \pi_k \frac{\Phi_{g+1,k}(t_{g+1}, \boldsymbol{y}_{-(g+1)})}{\Phi_{g+1,1}(t_{g+1}, \boldsymbol{y}_{-(g+1)})} = \sum_{l=1}^{L_{g+1}} \xi_l \frac{\Psi_{g+1,l}(t_{g+1}, \boldsymbol{y}_{-(g+1)})}{\Phi_{g+1,1}(t_{g+1}, \boldsymbol{y}_{-(g+1)})}$$
$$\Rightarrow \sum_{k=1}^{K_{g+1}} \pi_k f^{G-(g+1)}(\boldsymbol{y}_{-(g+1)}|\boldsymbol{\gamma}, \boldsymbol{\phi}_{-(g+1)}, \boldsymbol{p}_{-(g+1),k}) = \sum_{l=1}^{L_{g+1}} \xi_l f^{G-(g+1)}(\boldsymbol{y}_{-(g+1)}|\boldsymbol{\delta}, \boldsymbol{\psi}_{-(g+1)}, \boldsymbol{r}_{-(g+1),l}),$$

so Equation (**) holds for $G_0 = g+1$.

Consequently, by mathematical induction, we have shown that Equations (*) and (**) hold for any $G_0 \in \{1, \cdots, G\}$, which implies that $\boldsymbol{p}_1 = \boldsymbol{r}_1$ and $\boldsymbol{\phi} = \boldsymbol{\psi}$.

For $G_0 = G$ and $G_0 = G-1$, Equation (**) gives

$$\sum_{k=1}^{K_G} \pi_k = \sum_{l=1}^{L_G} \xi_l, \tag{16}$$
$$\sum_{k=1}^{K_{G-1}} \pi_k f^1(y_G|\boldsymbol{\gamma}, \phi_G, p_{Gk}) = \sum_{l=1}^{L_{G-1}} \xi_l f^1(y_G|\boldsymbol{\delta}, \phi_G, r_{Gl}). \tag{17}$$

For any two distinct elements $f^G(\boldsymbol{y}|\boldsymbol{\gamma}, \boldsymbol{\phi}, \boldsymbol{p}_1)$ and $f^G(\boldsymbol{y}|\boldsymbol{\gamma}, \boldsymbol{\phi}, \boldsymbol{p}_k), k = 2, 3, \cdots, K_{G-2}$, because there exist at least two different dimensions and $p_{g1} = p_{gk}$ with $g = 1, 2, \cdots, G-2$, $p_{G-1,k} \neq p_{G-1,1}$ and $p_{Gk} \neq p_{G1}$. Therefore, $K_G = K_{G-1} = 1$. Similarly, we have $L_G = L_{G-1} = 1$. Thus, Equation (16) and (17) turn to

$$\pi_1 = \xi_1$$
$$f^1(y_G|\boldsymbol{\gamma}, \phi_G, p_{G1}) = f^1(y_G|\boldsymbol{\delta}, \phi_G, r_{G1}), \forall y_G \in \mathbb{N}. \tag{18}$$

Plugging $y_G = 1$ and $y_G = 2$ into Equation (18), we have

$$\frac{1}{1+exp(\gamma_0+\gamma_1)} C_1^{\phi_G} p_{G1}(1-p_{G1})^{\phi_G} = \frac{1}{1+exp(\delta_0+\delta_1)} C_1^{\phi_G} p_{G1}(1-p_{G1})^{\phi_G}$$
$$\frac{1}{1+exp(\gamma_0+2\gamma_1)} C_2^{\phi_G+1} p_{G1}^2(1-p_{G1})^{\phi_G} = \frac{1}{1+exp(\delta_0+2\delta_1)} C_2^{\phi_G+1} p_{G1}^2(1-p_{G1})^{\phi_G},$$

therefore $\boldsymbol{\gamma} = \boldsymbol{\delta}$.

Plugging $\boldsymbol{\gamma} = \boldsymbol{\delta}, \boldsymbol{\phi} = \boldsymbol{\psi}, \boldsymbol{p}_1 = \boldsymbol{r}_1$ and $\pi_1 = \xi_1$ into Equation (5), we have

$$\sum_{k=2}^{K} \pi_k f^G(\boldsymbol{y}|\gamma, \boldsymbol{\phi}, \boldsymbol{p}_k) = \sum_{l=2}^{L} \xi_l f^G(\boldsymbol{y}|\gamma, \boldsymbol{\phi}, \boldsymbol{r}_l) \tag{19}$$

**Similarly, we can apply mathematical induction to prove that $\boldsymbol{p}_k = \boldsymbol{r}_k$ and $\pi_k = \xi_k$ sequentially for $k = 2, 3, \cdots, \min\{K, L\}$.** Finally, if $K \neq L$, without loss of generality, let us assume that $K > L$, then $\sum_{k=L+1}^{K} \pi_k = 1 - \sum_{k=1}^{L} \pi_k = 1 - \sum_{l=1}^{L} \xi_l = 0$, which contradicts with $\pi_k > 0$ for all $k = 1, 2, \cdots, K$. Thus, $K = L, \boldsymbol{\gamma} = \boldsymbol{\delta}, \boldsymbol{\phi} = \boldsymbol{\psi}, \boldsymbol{\pi} = \boldsymbol{\xi}$ and $\boldsymbol{p}_k = \boldsymbol{r}_k$ for all $k = 1, 2, \cdots, K$. Therefore, the class of all finite mixtures of $\mathcal{F}^G$ is identifiable. □

**Theorem 1.** *(The Complete Setting)*
*If $\pi_{bk} > 0$ for every batch $b$ and cell type $k$, given that (I) $\gamma_{b1} < 0$ for every $b$, (II) for any two cell types $k_1$ and $k_2$, there exist at least two differentially expressed genes $g_1$ and $g_2$— $\beta_{g_1 k_1} \neq \beta_{g_1 k_2}$ and $\beta_{g_2 k_1} \neq \beta_{g_2 k_2}$, and (III) for any two distinct cell-type pairs $(k_1, k_2) \neq (k_3, k_4)$, their differences in cell-type effects are not the same $\boldsymbol{\beta}_{k_1} - \boldsymbol{\beta}_{k_2} \neq \boldsymbol{\beta}_{k_3} - \boldsymbol{\beta}_{k_4}$, then BUSseq is identifiable (up to label switching) in the sense that $L_o(\boldsymbol{\Theta}|\boldsymbol{y}) = L_o(\boldsymbol{\Theta}^*|\boldsymbol{y})$ for any $\boldsymbol{y}$ implies that $\pi_{bk} = \pi^*_{b\rho(k)}, (\gamma_{b0}, \gamma_{b1}) = (\gamma^*_{b0}, \gamma^*_{b1}), \alpha_g + \beta_{gk} = \alpha^*_g + \beta^*_{g\rho(k)}, \nu_{gb} = \nu^*_{gb}, \delta_{bi} = \delta^*_{bi}$ and $\phi_{bg} = \phi^*_{bg}$ for every gene $g$ and batch $b$, where $\rho$ is a permutation of $\{1, 2, \cdots, K\}$.*

*Proof.* Let $\boldsymbol{Y}_b \in N^{n_b \times G}$ denote the data from batch $b$ and collect $\boldsymbol{Y} = \{\boldsymbol{Y}_b, 1 \leq b \leq B\}$ and $\boldsymbol{m}_{bik} = exp(\boldsymbol{\alpha} + \boldsymbol{\beta}_k + \boldsymbol{\nu}_b + \delta_{bi}\mathbf{1})$, then the marginal distribution for $f(\boldsymbol{Y}_b|\boldsymbol{\Theta}) = \prod_{i=1}^{n_b}[\sum_{k=1}^{K} \pi_{bk} f^G(\boldsymbol{y}_b|\boldsymbol{\gamma}_b, \boldsymbol{\phi}_b, \boldsymbol{m}_{bik})]$ with $f^G(\boldsymbol{y}_b|\boldsymbol{\gamma}_b, \boldsymbol{\phi}_b, \boldsymbol{m}_{bik}) \in \mathcal{F}^G$ reduces to a mixture of $G$-dimensional zero-inflated negative binomial (ZINB) model on batch $b$. Therefore, we can view the BUSseq model as a combination of $B$ ZINB models with the constraints that $\boldsymbol{\beta}_k^{(1)} = \cdots = \boldsymbol{\beta}_k^{(B)} = \boldsymbol{\beta}_k$ for each $k$ and $\boldsymbol{\alpha}^{(1)} = \cdots = \boldsymbol{\alpha}^{(B)} = \boldsymbol{\alpha}$.

According to conditions (I)-(III) and Lemma 1, the ZINB model for batch $b$ is identifiable up to label switching in the sense that $f(\boldsymbol{Y}_b|\boldsymbol{\Theta}) = f(\boldsymbol{Y}_b|\boldsymbol{\Theta}^*)$ for any $\boldsymbol{Y}_b$ implies that $\pi_{bk} = \pi^*_{b\rho_b(k)}, \boldsymbol{\gamma}_b = \boldsymbol{\gamma}^*_b, \boldsymbol{\alpha} + \boldsymbol{\beta}_k + \boldsymbol{\nu}_b + \delta_{bi}\mathbf{1} = log(\boldsymbol{m}_{bik}) = log(\boldsymbol{m}^*_{bi\rho_b(k)}) = \boldsymbol{\alpha}^* + \boldsymbol{\beta}^*_{\rho_b(k)} + \boldsymbol{\nu}^*_b + \delta^*_{bi}\mathbf{1}$ and $\boldsymbol{\phi}_b = \boldsymbol{\phi}^*_b$ for a permutation $\rho_b$ of $\{1, 2, \cdots, K\}$, where $\mathbf{1}$ denotes a vector of one with length $G$.

We first prove that the permutation $\rho_b$ is the same for all of the batches. Recall that we take the first cell type as the reference cell type with $\boldsymbol{\beta}_1 = 0$. Therefore, the ratio of mean expression levels between cell type $k$ and cell type one is

$$\frac{\boldsymbol{m}_{bik}}{\boldsymbol{m}_{bi1}} = \frac{\boldsymbol{m}^*_{bi\rho_b(k)}}{\boldsymbol{m}^*_{bi\rho_b(1)}} \Rightarrow exp(\boldsymbol{\beta}_k) = exp(\boldsymbol{\beta}^*_{\rho_b(k)} - \boldsymbol{\beta}^*_{\rho_b(1)}) \tag{20}$$

Notice the left hand side of Equation (20) is invariant to the batch indicator $b$, and therefore $\boldsymbol{\beta}^*_{\rho_b(k)} - \boldsymbol{\beta}^*_{\rho_b(1)} = \boldsymbol{\beta}^*_{\rho_1(k)} - \boldsymbol{\beta}^*_{\rho_1(1)}$ for every $k$. By condition (III), $\rho_b = \rho_1$ for every $b$.

Let us then compare $log(\boldsymbol{m}_{b1k})$ with $log(\boldsymbol{m}_{11k})$. Because $\boldsymbol{\nu}_1 = \boldsymbol{\nu}^*_1 = \mathbf{0}$ and $\delta_{b1} = \delta^*_{b1} = 0$, we have

$$\boldsymbol{\alpha} + \boldsymbol{\beta}_k = \boldsymbol{\alpha}^* + \boldsymbol{\beta}^*_{\rho(k)}, \boldsymbol{\alpha} + \boldsymbol{\beta}_k + \boldsymbol{\nu}_b = \boldsymbol{\alpha}^* + \boldsymbol{\beta}^*_{\rho(k)} + \boldsymbol{\nu}^*_b. \tag{21}$$

Thus, we have proven $\boldsymbol{\nu}_b = \boldsymbol{\nu}_b^*$.

Next we compare $log(\boldsymbol{m}_{bik})$ with $log(\boldsymbol{m}_{b1k})$ for each batch. Then, we have

$$\boldsymbol{\alpha} + \boldsymbol{\beta}_k + \boldsymbol{\nu}_b = \boldsymbol{\alpha}^* + \boldsymbol{\beta}^*_{\rho(k)} + \boldsymbol{\nu}_b, \boldsymbol{\alpha} + \boldsymbol{\beta}_k + \boldsymbol{\nu}_b + \delta_{bi}\mathbf{1} = \boldsymbol{\alpha}^* + \boldsymbol{\beta}^*_{\rho(k)} + \boldsymbol{\nu}_b + \delta^*_{bi}\mathbf{1}. \tag{22}$$

Consequently, $\delta_{bi} = \delta^*_{bi}$ for any cell $i$ in any batch. Therefore, BUSseq is identifiable (up to label switching). □

**Theorem 2.** *(The Reference Panel Design)*
*If there are a total of $K$ cell types $\cup_{b=1}^B C_b = \{1, 2, \cdots, K\}$, where $C_b$ denotes the cell types that are present in batch $b$, the number of cell types existing in batch $b$ $K_b = |C_b| \geq 2$ for every batch $b$, and there exists a batch $\tilde{b}$ such that it contains all of the cell types $C_{\tilde{b}} = \{1, 2, \cdots, K\}$, then given that conditions (I)-(III) hold, BUSseq is identifiable (up to label switching).*

*Proof.* In the reference panel design, any batch $b$ shares at least two cell types with the first batch. If we compare the two distinct cell types $k_1$ and $k_2$ shared by batch $b$ and batch one in terms of the log-scale mean expression levels, respectively, then we have

$$\left.\begin{aligned} \boldsymbol{\alpha} + \boldsymbol{\beta}_{k_1} + \boldsymbol{\nu}_b + \delta_{bi}\mathbf{1} = \boldsymbol{\alpha}^* + \boldsymbol{\beta}^*_{\rho_b(k_1)} + \boldsymbol{\nu}^*_b + \delta^*_{bi}\mathbf{1}, \\ \boldsymbol{\alpha} + \boldsymbol{\beta}_{k_2} + \boldsymbol{\nu}_b + \delta_{bi}\mathbf{1} = \boldsymbol{\alpha}^* + \boldsymbol{\beta}^*_{\rho_b(k_2)} + \boldsymbol{\nu}^*_b + \delta^*_{bi}\mathbf{1}, \end{aligned}\right\} \Rightarrow \boldsymbol{\beta}_{k_1} - \boldsymbol{\beta}_{k_2} = \boldsymbol{\beta}^*_{\rho_b(k_1)} - \boldsymbol{\beta}^*_{\rho_b(k_2)},$$

$$\left.\begin{aligned} \boldsymbol{\alpha} + \boldsymbol{\beta}_{k_1} + \delta_{1i}\mathbf{1} = \boldsymbol{\alpha}^* + \boldsymbol{\beta}^*_{\rho_1(k_1)} + \delta^*_{1i}\mathbf{1}, \\ \boldsymbol{\alpha} + \boldsymbol{\beta}_{k_2} + \delta_{1i}\mathbf{1} = \boldsymbol{\alpha}^* + \boldsymbol{\beta}^*_{\rho_1(k_2)} + \delta^*_{1i}\mathbf{1}. \end{aligned}\right\} \Rightarrow \boldsymbol{\beta}_{k_1} - \boldsymbol{\beta}_{k_2} = \boldsymbol{\beta}^*_{\rho_1(k_1)} - \boldsymbol{\beta}^*_{\rho_1(k_2)}.$$

Further, according to condition (III), $\boldsymbol{\beta}_{k_1} - \boldsymbol{\beta}_{k_2} = \boldsymbol{\beta}^*_{\rho_b(k_1)} - \boldsymbol{\beta}^*_{\rho_b(k_2)} = \boldsymbol{\beta}^*_{\rho_1(k_1)} - \boldsymbol{\beta}^*_{\rho_1(k_2)}$ implies that $\rho_b(k) = \rho_1(k)$ for each cell type $k \in C_b$ $(b \geq 2)$.

Finally, similar to Equations (21) and (22), for a shared cell type $k$ between batch $b$ and batch one, we have

$$\begin{aligned} \boldsymbol{\alpha} + \boldsymbol{\beta}_k &= \boldsymbol{\alpha}^* + \boldsymbol{\beta}^*_{\rho_1(k)} \\ \boldsymbol{\alpha} + \boldsymbol{\beta}_k + \boldsymbol{\nu}_b &= \boldsymbol{\alpha}^* + \boldsymbol{\beta}^*_{\rho_b(k)} + \boldsymbol{\nu}^*_b \\ \boldsymbol{\alpha} + \boldsymbol{\beta}_k + \boldsymbol{\nu}_b + \delta_{bi}\mathbf{1} &= \boldsymbol{\alpha}^* + \boldsymbol{\beta}^*_{\rho_b(k)} + \boldsymbol{\nu}^*_b + \delta^*_{bi}\mathbf{1}. \end{aligned}$$

Thus, $\rho_b(k) = \rho_1(k)$ for each $k \in C_b$ $(b \geq 2)$ implies that $\boldsymbol{\nu}_b = \boldsymbol{\nu}^*_b$ and $\delta_{bi} = \delta^*_{bi}$. □

**Theorem 3.** *(The Chain-type Design)*
*If there are a total of $K$ cell types $\cup_{b=1}^B C_b = \{1, 2, \cdots, K\}$ and every two consecutive batches share at least two cell types $|C_b \cap C_{b-1}| \geq 2$ for all $b \geq 2$, then given that conditions (I)-(III) hold, BUSseq is identifiable (up to label switching).*

*Proof.* Our objective is to prove that for any two distinct batches $b$ and $\tilde{b}$, $\rho_b(k) = \rho_{\tilde{b}}(k)$ holds for any cell type $k \in C_b \bigcap C_{\tilde{b}}$ shared by these two batches.

First, we prove that $\rho_b(k) = \rho_{b-1}(k)$ for the shared cell types $k \in C_b \bigcap C_{b-1}, 2 \leq b \leq B$ in any two consecutive batches. Notice that $|C_b \cap C_{b-1}| \geq 2$, so for any two shared cell types $k_1$ and $k_2$ between batch $b$ and batch $b-1$, we have

$$\left.\begin{array}{l}\boldsymbol{\alpha} + \boldsymbol{\beta}_{k_1} + \boldsymbol{\nu}_b + \delta_{bi}\mathbf{1} = \boldsymbol{\alpha}^* + \boldsymbol{\beta}^*_{\rho_b(k_1)} + \boldsymbol{\nu}^*_b + \delta^*_{bi}\mathbf{1},\\ \boldsymbol{\alpha} + \boldsymbol{\beta}_{k_2} + \boldsymbol{\nu}_b + \delta_{bi}\mathbf{1} = \boldsymbol{\alpha}^* + \boldsymbol{\beta}^*_{\rho_b(k_2)} + \boldsymbol{\nu}^*_b + \delta^*_{bi}\mathbf{1},\end{array}\right\} \Rightarrow \boldsymbol{\beta}_{k_1} - \boldsymbol{\beta}_{k_2} = \boldsymbol{\beta}^*_{\rho_b(k_1)} - \boldsymbol{\beta}^*_{\rho_b(k_2)}$$

$$\left.\begin{array}{l}\boldsymbol{\alpha} + \boldsymbol{\beta}_{k_1} + \boldsymbol{\nu}_{b-1} + \delta_{b-1,i}\mathbf{1} = \boldsymbol{\alpha}^* + \boldsymbol{\beta}^*_{\rho_{b-1}(k_1)} + \boldsymbol{\nu}^*_{b-1} + \delta^*_{b-1,i}\mathbf{1},\\ \boldsymbol{\alpha} + \boldsymbol{\beta}_{k_2} + \boldsymbol{\nu}_{b-1} + \delta_{b-1,i}\mathbf{1} = \boldsymbol{\alpha}^* + \boldsymbol{\beta}^*_{\rho_{b-1}(k_2)} + \boldsymbol{\nu}^*_{b-1} + \delta^*_{b-1,i}\mathbf{1}.\end{array}\right\} \Rightarrow \boldsymbol{\beta}_{k_1} - \boldsymbol{\beta}_{k_2} = \boldsymbol{\beta}^*_{\rho_{b-1}(k_1)} - \boldsymbol{\beta}^*_{\rho_{b-1}(k_2)}$$

Further, according to condition (III), $\boldsymbol{\beta}_{k_1} - \boldsymbol{\beta}_{k_2} = \boldsymbol{\beta}^*_{\rho_b(k_1)} - \boldsymbol{\beta}^*_{\rho_b(k_2)} = \boldsymbol{\beta}^*_{\rho_{b-1}(k_1)} - \boldsymbol{\beta}^*_{\rho_{b-1}(k_2)}$ implies that $\rho_b(k) = \rho_{b-1}(k)$ for $2 \leq b \leq B, k \in C_b \bigcap C_{b-1}$.

Consequently, for a cell type $k \in C_b \bigcap C_{b-1}$ shared by two consecutive batches $b$ and $b-1$, similar to Equation (21), we have

$$\left.\begin{array}{l}\boldsymbol{\alpha} + \boldsymbol{\beta}_k + \boldsymbol{\nu}_{b-1} = \boldsymbol{\alpha}^* + \boldsymbol{\beta}^*_{\rho_{b-1}(k)} + \boldsymbol{\nu}^*_{b-1}\\ \boldsymbol{\alpha} + \boldsymbol{\beta}_k + \boldsymbol{\nu}_b = \boldsymbol{\alpha}^* + \boldsymbol{\beta}^*_{\rho_b(k)} + \boldsymbol{\nu}^*_b\end{array}\right\} \Rightarrow \boldsymbol{\nu}_b - \boldsymbol{\nu}_{b-1} = \boldsymbol{\nu}^*_b - \boldsymbol{\nu}^*_{b-1}.$$

Because $\boldsymbol{\nu}_1 = \boldsymbol{\nu}^*_1 = 0$, $\boldsymbol{\nu}_b = \sum_{j=2}^b (\boldsymbol{\nu}_j - \boldsymbol{\nu}_{j-1}) = \sum_{j=2}^b (\boldsymbol{\nu}^*_j - \boldsymbol{\nu}^*_{j-1}) = \boldsymbol{\nu}^*_b$. Moreover, similar to Equation (22), we have $\delta_{bi} = \delta^*_{bi}$ for each cell $i$ of each batch $b$.

Now for any two distinct batches $b$ and $\tilde{b}$, we can directly compare the mean expression levels of their shared cell type $k \in C_b \bigcap C_{\tilde{b}}$:

$$\left.\begin{array}{l}\boldsymbol{\alpha} + \boldsymbol{\beta}_k + \boldsymbol{\nu}_b + \delta_{bi}\mathbf{1} = \boldsymbol{\alpha}^* + \boldsymbol{\beta}^*_{\rho_b(k)} + \boldsymbol{\nu}_b + \delta_{bi}\mathbf{1}\\ \boldsymbol{\alpha} + \boldsymbol{\beta}_k + \boldsymbol{\nu}_{\tilde{b}} + \delta_{\tilde{b}i}\mathbf{1} = \boldsymbol{\alpha}^* + \boldsymbol{\beta}^*_{\rho_{\tilde{b}}(k)} + \boldsymbol{\nu}_{\tilde{b}} + \delta_{\tilde{b}i}\mathbf{1}\end{array}\right\} \Rightarrow \boldsymbol{\beta}^*_{\rho_b(k)} = \boldsymbol{\beta}^*_{\rho_{\tilde{b}}(k)}$$

□

**Theorem 4.** *(The Connected Design)*
*We define a batch graph $G = (V, E)$. Each node $b \in V$ represents a batch. There is an edge $e \in E$ between two nodes $b_1$ and $b_2$ if and only if batches $b_1$ and $b_2$ share at least two cell types. If the batch graph is connected and conditions (I)-(III) hold, then BUSseq is identifiable (up to label switching).*

*Proof.* Our object is to prove that for any two distinct batches $b$ and $\tilde{b}$, $\rho_b(k) = \rho_{\tilde{b}}(k)$ holds for any cell type $k \in C_b \bigcap C_{\tilde{b}}$ shared by these two batches. At the same time, $\boldsymbol{\nu}_b = \boldsymbol{\nu}^*_b$ and $\delta_{bi} = \delta^*_{bi}$ for each cell $i$ in batch $b$.

For any two connected batches $(b_1, b_2)$, we have $|C_{b_1} \cap C_{b_2}| \geq 2$. Thus, for any two shared cell types $k_1, k_2 \in C_{b_1} \cap C_{b_2}$, we have

$$\left.\begin{array}{l}\boldsymbol{\alpha}+\boldsymbol{\beta}_{k_1}+\boldsymbol{\nu}_{b_1}+\delta_{b_1,i}\mathbf{1}=\boldsymbol{\alpha}^*+\boldsymbol{\beta}^*_{\rho_{b_1}(k_1)}+\boldsymbol{\nu}^*_{b_1}+\delta^*_{b_1,i}\mathbf{1},\\ \boldsymbol{\alpha}+\boldsymbol{\beta}_{k_2}+\boldsymbol{\nu}_{b_1}+\delta_{b_1,i}\mathbf{1}=\boldsymbol{\alpha}^*+\boldsymbol{\beta}^*_{\rho_{b_1}(k_2)}+\boldsymbol{\nu}^*_{b_1}+\delta^*_{b_1,i}\mathbf{1},\end{array}\right\}\Rightarrow \boldsymbol{\beta}_{k_1}-\boldsymbol{\beta}_{k_2}=\boldsymbol{\beta}^*_{\rho_{b_1}(k_1)}-\boldsymbol{\beta}^*_{\rho_{b_1}(k_2)}$$

$$\left.\begin{array}{l}\boldsymbol{\alpha}+\boldsymbol{\beta}_{k_1}+\boldsymbol{\nu}_{b_2}+\delta_{b_2,i}\mathbf{1}=\boldsymbol{\alpha}^*+\boldsymbol{\beta}^*_{\rho_{b_2}(k_1)}+\boldsymbol{\nu}^*_{b_2}+\delta^*_{b_2,i}\mathbf{1},\\ \boldsymbol{\alpha}+\boldsymbol{\beta}_{k_2}+\boldsymbol{\nu}_{b_2}+\delta_{b_2,i}\mathbf{1}=\boldsymbol{\alpha}^*+\boldsymbol{\beta}^*_{\rho_{b_2}(k_2)}+\boldsymbol{\nu}^*_{b_2}+\delta^*_{b_2,i}\mathbf{1}.\end{array}\right\}\Rightarrow \boldsymbol{\beta}_{k_1}-\boldsymbol{\beta}_{k_2}=\boldsymbol{\beta}^*_{\rho_{b_2}(k_1)}-\boldsymbol{\beta}^*_{\rho_{b_2}(k_2)}$$

Further, according to condition (III), $\boldsymbol{\beta}_{k_1}-\boldsymbol{\beta}_{k_2}=\boldsymbol{\beta}^*_{\rho_{b_1}(k_1)}-\boldsymbol{\beta}^*_{\rho_{b_1}(k_2)}=\boldsymbol{\beta}^*_{\rho_{b_2}(k_1)}-\boldsymbol{\beta}^*_{\rho_{b_2}(k_2)}$ implies that $\rho_{b_1}(k)=\rho_{b_2}(k)$ for $k \in C_{b_1}\bigcap C_{b_2}$.

Consequently, for a cell type $k \in C_{b_1}\bigcap C_{b_2}$ shared by two connected batches $b_1$ and $b_2$, similar to Equation (21), we have

$$\left.\begin{array}{l}\boldsymbol{\alpha}+\boldsymbol{\beta}_{k}+\boldsymbol{\nu}_{b_1}=\boldsymbol{\alpha}^*+\boldsymbol{\beta}^*_{\rho_{b_1}(k)}+\boldsymbol{\nu}^*_{b_1}\\ \boldsymbol{\alpha}+\boldsymbol{\beta}_{k}+\boldsymbol{\nu}_{b_2}=\boldsymbol{\alpha}^*+\boldsymbol{\beta}^*_{\rho_{b_2}(k)}+\boldsymbol{\nu}^*_{b_2}\end{array}\right\}\Rightarrow \boldsymbol{\nu}_{b_1}-\boldsymbol{\nu}^*_{b_1}=\boldsymbol{\nu}_{b_2}-\boldsymbol{\nu}^*_{b_2}.$$

Because of the connectivity of the batch graph $G$, we can find a path $(1, b_1, b_2, \cdots, b_k, b), k \leq B-2$ between any batch $b$ and the first batch in the batch graph $G$ such that $\boldsymbol{\nu}_b-\boldsymbol{\nu}^*_b=\boldsymbol{\nu}_{b_k}-\boldsymbol{\nu}^*_{b_k}=\cdots=\boldsymbol{\nu}_{b_1}-\boldsymbol{\nu}^*_{b_1}=\boldsymbol{\nu}_1-\boldsymbol{\nu}^*_1$. Notice that $\boldsymbol{\nu}_1=\boldsymbol{\nu}^*_1=0$, so we have $\boldsymbol{\nu}_b=\boldsymbol{\nu}^*_b$. Moreover, similar to Equation (22), we have $\delta_{bi}=\delta^*_{bi}$ for each cell $i$ in the batch $b$.

Now for any two distinct batches $b$ and $\tilde{b}$, we can directly compare the mean expression levels of their shared cell type $k \in C_b \bigcap C_{\tilde{b}}$:

$$\left.\begin{array}{l}\boldsymbol{\alpha}+\boldsymbol{\beta}_{k}+\boldsymbol{\nu}_{b}+\delta_{bi}\mathbf{1}=\boldsymbol{\alpha}^*+\boldsymbol{\beta}^*_{\rho_{b}(k)}+\boldsymbol{\nu}_{b}+\delta_{bi}\mathbf{1}\\ \boldsymbol{\alpha}+\boldsymbol{\beta}_{k}+\boldsymbol{\nu}_{\tilde{b}}+\delta_{\tilde{b}i}\mathbf{1}=\boldsymbol{\alpha}^*+\boldsymbol{\beta}^*_{\rho_{\tilde{b}}(k)}+\boldsymbol{\nu}_{\tilde{b}}+\delta_{\tilde{b}i}\mathbf{1}\end{array}\right\}\Rightarrow \boldsymbol{\beta}^*_{\rho_b(k)}=\boldsymbol{\beta}^*_{\rho_{\tilde{b}}(k)},$$

which implies that $\rho_b(k)=\rho_{\tilde{b}}(k)$ according to condition (II). □
